# Dissecting unsupervised learning through hidden Markov modelling in electrophysiological data

**DOI:** 10.1101/2023.01.19.524547

**Authors:** Laura Masaracchia, Felipe Fredes, Mark W. Woolrich, Diego Vidaurre

## Abstract

Unsupervised, data-driven methods are commonly used in neuroscience to automatically decompose data into interpretable patterns. These patterns differ from one another depending on the assumptions of the models. How these assumptions affect specific data decompositions in practice, however, is often unclear, which hinders model applicability and interpretability. For instance, the hidden Markov model (HMM) automatically detects characteristic, recurring activity patterns (so-called *states*) from time series data. States are defined by a certain probability distribution, whose state-specific parameters are estimated from the data. But what specific features, from all of those that the data contain, do the states capture? That depends on the choice of probability distribution and on other model hyperparameters. Using both synthetic and real data, we aim at better characterizing the behavior of two HMM types that can be applied to electrophysiological data. Specifically, we study which differences in data features (such as frequency, amplitude or signal-to-noise ratio) are more salient to the models and therefore more likely to drive the state decomposition. Overall, we aim at providing guidance for an appropriate use of this type of analysis on one or two-channel neural electrophysiological data, and an informed interpretation of its results given the characteristics of the data and the purpose of the analysis.

**NEW & NOTEWORTHY:** Compared to classical supervised methods, unsupervised methods of analysis have the advantage to be freer of subjective biases. However, it is not always clear what aspects of the data these methods are most sensitive to, which complicates interpretation. Focusing on the Hidden Markov Model, commonly used to describe electrophysiological data, we explore in detail the nature of its estimates through simulations and real data examples, providing important insights about what to expect from these models.

## 1. INTRODUCTION

In empirical neuroscience, we often use supervised methods of analysis to investigate the mechanistic underpinnings of cognitive processing. These are supervised in the sense that they assume a certain preconception of the patterns of interest, and search for their expressions in the data accordingly. Some examples are decoding analysis [1, 2, 3], analyses of evoked response potentials (ERP) [4, 5, 6] and methods to characterize oscillations [7, 8, 9]. Overall, supervised approaches rely on prior knowledge and definitions of the features of interest, which could potentially be incomplete or imprecise.

An alternative approach is the use of data-driven, unsupervised models to automatically extract patterns from brain data without a prior definition of such patterns. For example, a clustering algorithm run on the time series data would automatically find a collection of patterns that was not defined beforehand, and it is then the researcher’s task to determine what these patterns might mean neurobiologically. While unsupervised methods are potentially less biased and therefore can have a better chance of finding new information, they may also be more difficult to interpret. For this reason, it is important to, at least, have a clear understanding of what elements in the data are more salient for the chosen unsupervised algorithm, in order to elucidate what a given decomposition is really capturing.

We focus on the Hidden Markov Model (HMM) [10], that have been used to investigate various domains, such as resting state dynamics in wakefulness [11, 12, 13, 14, 15] and sleep [16], perceptual processing [17], memory replay [18] and higher-order cognition [19]. These studies are examples of how the HMM can automatically identify patterns of activity (so-called *states*) without the need of providing beforehand a stringent, explicit definition of what would constitute a meaningful pattern. But what aspects of the data are more salient to the model -i.e., what is really driving the state inference? That depends on the choice of the observation model, which is the probability distribution used to represent the states. Two different observation models might have different views of saliency (meaning that different modulations in the data are considered more important for different models), and therefore behave differently. Often, the choice of one model over another is left to trial and error. While empirical comparisons have been made in the past between different observation models for specific purposes [20], no study has rigorously investigated what specific features within the data are the different state definitions most sensitive to. We reason that a better understanding of the various alternatives is important to fully leverage the benefits of this type of unsupervised analysis and interpret their outputs. Here, focusing on raw, low-dimensional (one or two channels) electrophysiological data, we explore two types of HMM: the HMM-MAR, where the states are multivariate autoregressive models [21]; and the HMM-TDE (time-delay embedded), where the states correspond to autocovariance patterns in the signal [12].

We first use synthetic data to investigate how sensitive the two observation models are to variations in frequency, amplitude, signal-to-noise ratio, and amount of data. Then, we analyze how these characteristics affect the estimation of the states in the HMMs. Finally, we examine the behavior of the models on two different real data modalities, LFP from mice in resting state and MEG from humans performing a simple motor task. In summary, our results show what to expect from the HMM states according to data conditions and analysis settings. On these grounds, we provide some recommendations on which model to use depending on the data and the purpose of the analysis.

## 2. MATERIALS & METHODS

Using both real and simulated electrophysiological data, this study investigated the behavior of two varieties of the HMM: the HMM-MAR [21] and the HMM-TDE [12]. First, a sensitivity analysis on the observation models (MAR and TDE) was conducted, with respect to different data features and model hyperparameters. Secondly, the full HMM behavior was assessed, using both synthetic and real electrophysiological data: LFP data from a mouse hippocampus (during wakeful resting) and MEG data of human subjects performing a simple button press task. A schematic of the study is shown in **Figure 1**.

**Figure 1:**
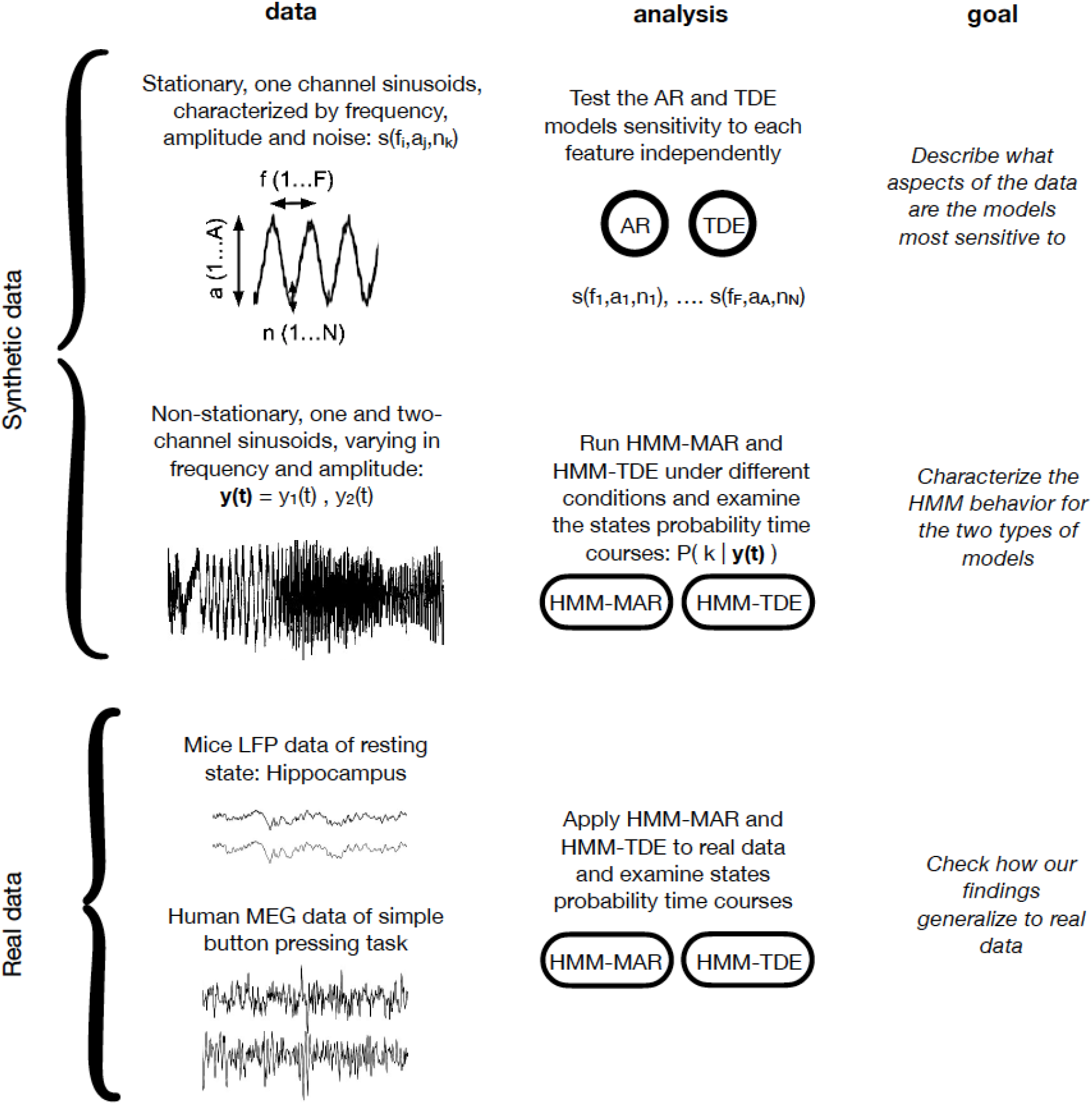
Scheme of the workflow, including data, analysis type and goals of each block.

### 2.1 Data

#### 2.1.1 Synthetic data

##### Synthetic, stationary, one-channel signals

We tested the stand-alone observation model sensitivity to different data features using single-sinusoidal (stationary) signals, defined by frequency *f*, sampling frequency *F*, signal length *T*, amplitude *a*, plus some random Gaussian noise, parametrized by its variance *v*. While *F* was kept constant, a range of values was established for the other features, and stationary sinusoids were sampled with all the combinations of the different features’ values. Specifically, *f* ranged from 0.5 to 45 Hz in steps of 0.02 Hz, *a* ranged between 0.5 and 10.5 (arbitrary units) in steps of 0.5 for the MAR model, and between 0.5 and 25 in multiplicative steps (proportions of 5.0) for the TDE model, *v* was either 0.5 or 1.0, and *T* was 2, 5, 10 or 15 seconds. *F* was 250 Hz in all cases (see **Figure 2a** for some signals examples).

**Figure 2:**
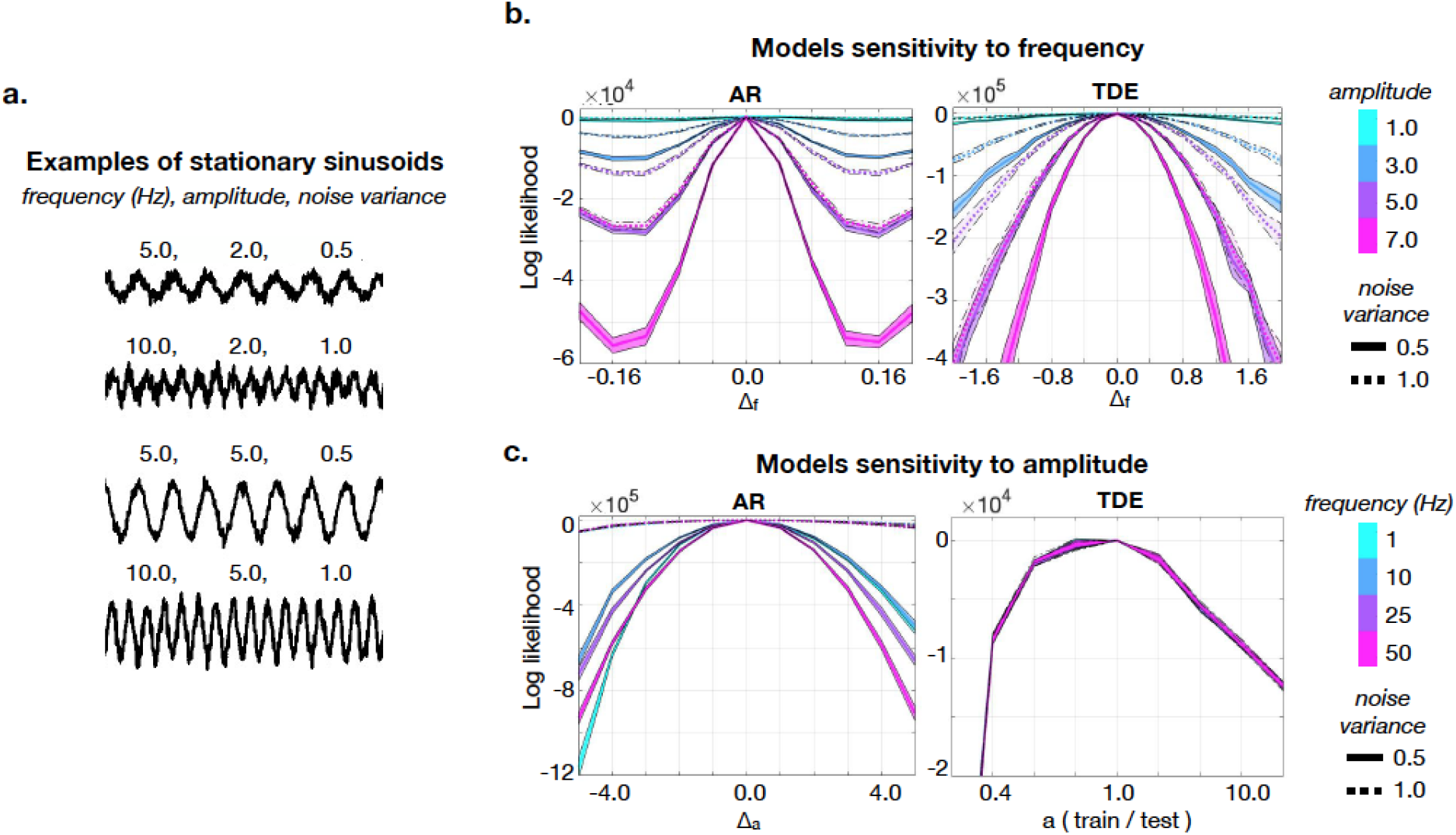
Analysis on the observation models. **a**. Example of signals used (stationary sinusoids), each defined by frequency, amplitude and noise variance. **b**. The plots show how the AR (left) and the TDE (right) models can tell apart two signals that differ only in frequency by an amount of Δ_*f*_ Hz (test frequency minus training frequency), for different values of their amplitude and of their noise content. The measure used is the logarithm of the likelihood ratio between train and test signal (given a fixed test signal and models trained on many training signals). Note the different scale of the x-axis for AR and TDE, indicating the different frequency sensitivity of the two models. Each solid line of the plot represents analyses for noise variance equal to 0.5, and each dotted line corresponds to noise variance equal to 1.0. By manipulating amplitude and noise variance, the plots show how the models perform for different signal to noise ratio (SNR) values. Here, AR order *P*=3, and TDE lags *L*=21, in steps of *S*=1; signal length *T*=10 seconds (25000 data points) **c**. AR and TDE sensitivity to amplitude, expressed as Δ_*a*_ for the AR model (training amplitude minus test amplitude) and as training amplitude in proportion to the target test amplitude for the TDE model (once again, signaling the different sensitivity to amplitude of the two models), for different values of frequency and of noise variance. Order, lags and signal length set as in **b**.

##### Synthetic, non-stationary, one-channel signals, with one single frequency

We used synthetic, non-stationary, one-channel signals to explore how the HMM-MAR and HMM-TDE segmented the time series into states visits (i.e., how they defined states and assigned a state probability per data point). The non-stationary oscillatory signals were created in such a way that signals had time varying frequency (ranging between 0.1 and 45.0 Hz) and amplitude (ranging between 0.1 and 10.0). Note that this generative model is more general (i.e., with more degrees of freedom) than the HMM, which assumes signals with quasi-stationary periods of sustained oscillations. Specifically, the instantaneous frequency *f(t)* and instantaneous amplitude *a(t)* were generated as two independent random walks, bounded within the chosen frequency and amplitude ranges, and where the step size at each time point was drawn from a normal distribution. The non-stationary, oscillatory signals were synthesized as a unique session of *T*=50000 data points, with sampling frequency *F*=250 Hz, and noise variance *v=*1.0 (see **Figure 3a** for an example), and then fed to the HMMs. The simulated session was then divided into *N*=100 trials of *t*_*N*_=500 timepoints each, to ease a cross-validation scheme for our prediction analysis (see **Section 2.3.2** for details).

**Figure 3:**
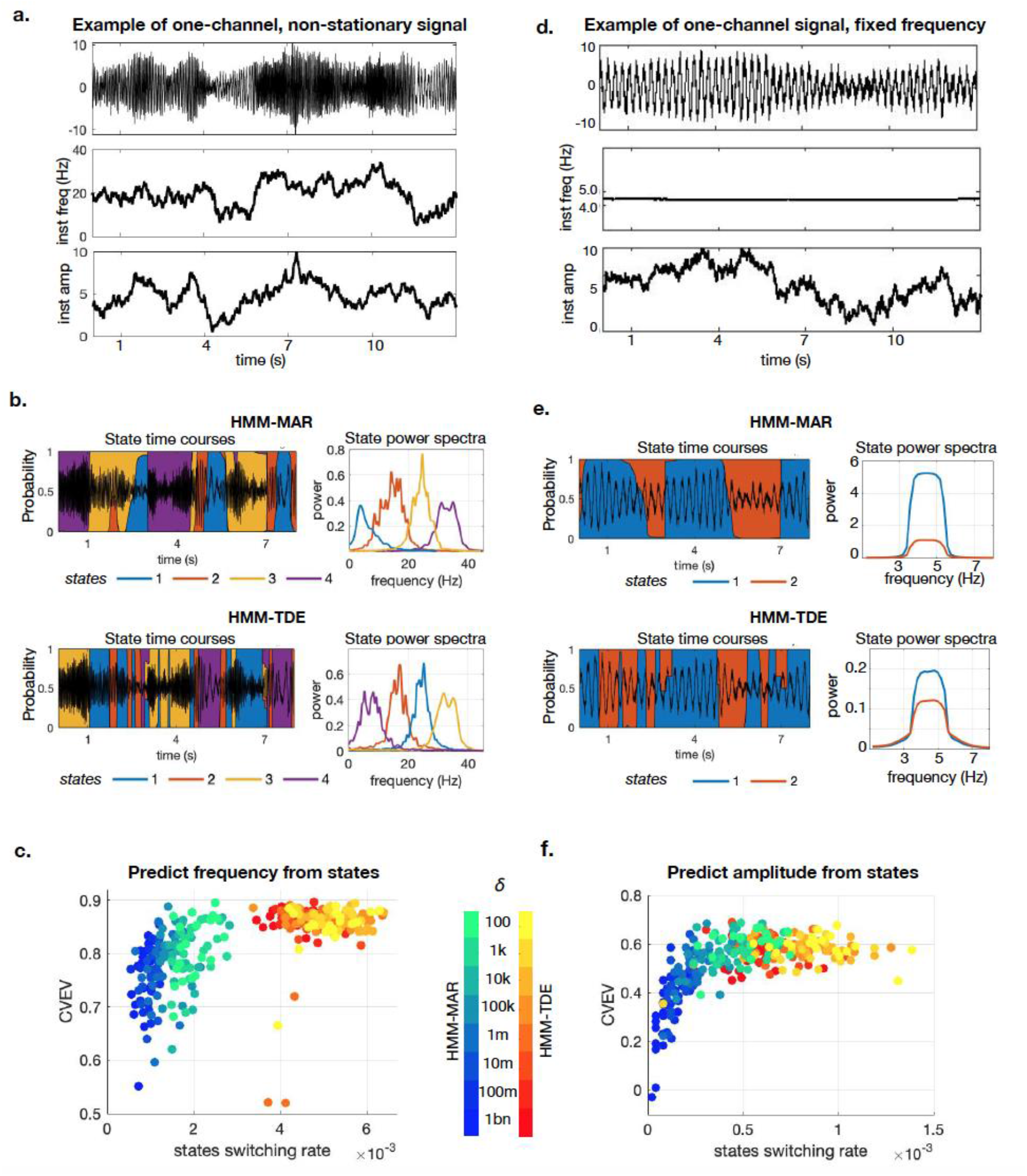
HMM on one-channel non-stationary data with one single frequency. **a**. Example of signal varying in frequency (instantaneous frequency shown in the middle panel) and amplitude (instantaneous amplitude in the bottom panel). **b**. Example of the probabilistic state time courses and state power spectra of HMM-MAR and HMM-TDE applied to the signal in **a**. Here, transition probability matrix prior *δ* = 10k, HMM-MAR order *P*=3, HMM-TDE lags *L*=15 (in steps of *S*=3). **c**. Cross validated explained variance of the HMM states predicting the ground truth frequency of non-stationary signals (like in a), for 10 repetitions of the experiment, for different values of the average state switching rate, manipulated via *δ* (order and lags set as in **b**). **d**. Similarly to **a**: example of a synthetic signal varying mostly in amplitude. **e**. Example of probabilistic state time courses and state power spectra of HMM-MAR and HMM-TDE applied to the signal in **d**. Here, *δ* = 10k, *P*=3, *L* = 15, *S*=3. **f**. Cross validated explained variance of the HMM states predicting the ground truth amplitude of the signals as a function of the average state switching rate (varying *δ*, order and lags set as in **e**), for 10 repetitions of the experiment.

##### Synthetic, non-stationary, one-channel signals, with two coexisting frequencies

The HMMs were also tested on one-channel, synthetic signals with two coexisting frequency oscillations, one in a lower frequency band and one in a higher frequency band. The relative power of the two frequency bands was modulated such that the lower frequency component had a larger power than the higher frequency by a certain factor. Specifically, the signal *x* was generated from two independent, non-stationary, one-channel signals (each with a single frequency content, but in a different frequency band) as:

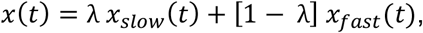

where *x*_*slow*_ had instantaneous frequency between 1 and 15 Hz, and *x*_*fast*_ had instantaneous frequency between 20 and 34 Hz. The coefficient *λ*, ranging between 0.8 and 0.5, modulated their power ratio, so that the signal in the lower frequency band (*x*_*slow*_) had higher or equal power than the signal in the higher frequency band (*x*_*fast*_). The data were generated as a continuous session of *T*=50000 points (see signal example in **Figure 4a**), fed to the HMMs and then reshaped into *N*=100 trials of *t*_*N*_=500 points for the prediction analyses (see below). Note that here, because of the presence of two coexisting frequencies, the MAR order *P* was increased to 7, so that it could capture more than one frequency [22].

**Figure 4:**
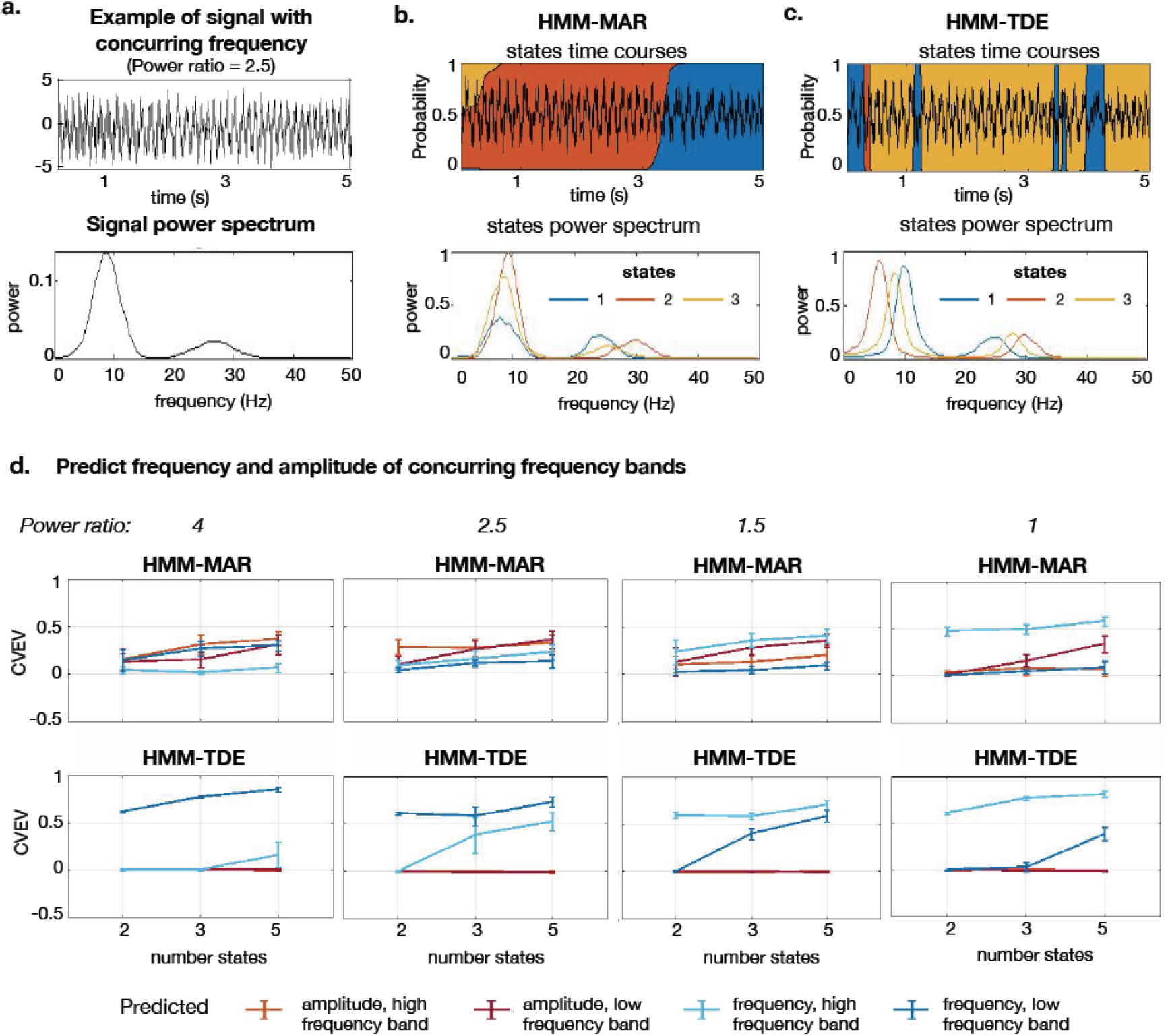
HMM-MAR and HMM-TDE applied to non-stationary, one-channel data with two coexisting frequencies. **a**. Example of signal with two concurring frequency contributions (power ratio = 2.5, upper panel) and its power spectrum (bottom panel). **b**. Example of states time courses (upper panel) and states power spectrum (bottom panel) of HMM-MAR applied to the signal in **a**. Here HMM-MAR order P=7, *δ*=100k. **c**. Example of states time courses (upper panel) and states power spectrum (bottom panel) of HMM-TDE applied to the signal in **a**. Here HMM-TDE lags L=15, spaced by S=3, *δ*=100k. **d**. Average Cross Validated Explained Variance (CVEV) and standard deviation (over 10 repetitions) of the predicted frequency and amplitude of the different frequency contributions (low and high frequency band) from the HMM states time courses, as a function of the states number K and for different power ratio scenarios. For all the analyses, we set *δ*=100k, HMM-MAR *P*=7, HMM-TDE *L*=15, with *S*=3.

##### Synthetic, non-stationary, two-channel signals

The HMMs were then tested on two-channel data showing time-varying between-channel coherence (to simulate simplified functional connectivity in the data). In particular, the two periodically coherent channels, *x*_1_ and *x*_2_, were generated combining three independent, non-stationary, one-channel signals (sampled as detailed above) *a, b* and *c*, as follows:

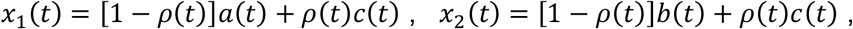

where *ρ(t)* modulates the channels similarity at each time point, and was generated as a smooth square wave (between 0 and 1) with a 4 seconds (1000 points) period. This way, when *ρ(t) = 1* the two channels were equal to *c(t)* and therefore maximally coherent; however, when *ρ(t) = 0*, the two channels corresponded, respectively, to *a(t)* and *b(t)*, which were independently generated —but, crucially, not forced to be strictly uncorrelated, and hence they could still exhibit some residual correlation due to sampling variability. In this sense, *ρ(t)* was not a real measure for coherence, and we adopted instead the two channels’ instantaneous empirical correlation *r(t)*, computed within a sliding window, as a surrogate measure of the ground-truth channels’ coherence. The sliding window’s size was chosen such that *r(t)* matched best *ρ(t)* = 1, i.e., when the two channels were actually the same signal, and their ground-truth coherence was known. Once again, the data were generated as a continuous session of *T*=50000 points (see example in **Figure 5a**), fed to the HMMs and then reshaped into *N*=100 trials of *t*_*N*_=500 points for the prediction analysis (see below).

**Figure 5:**
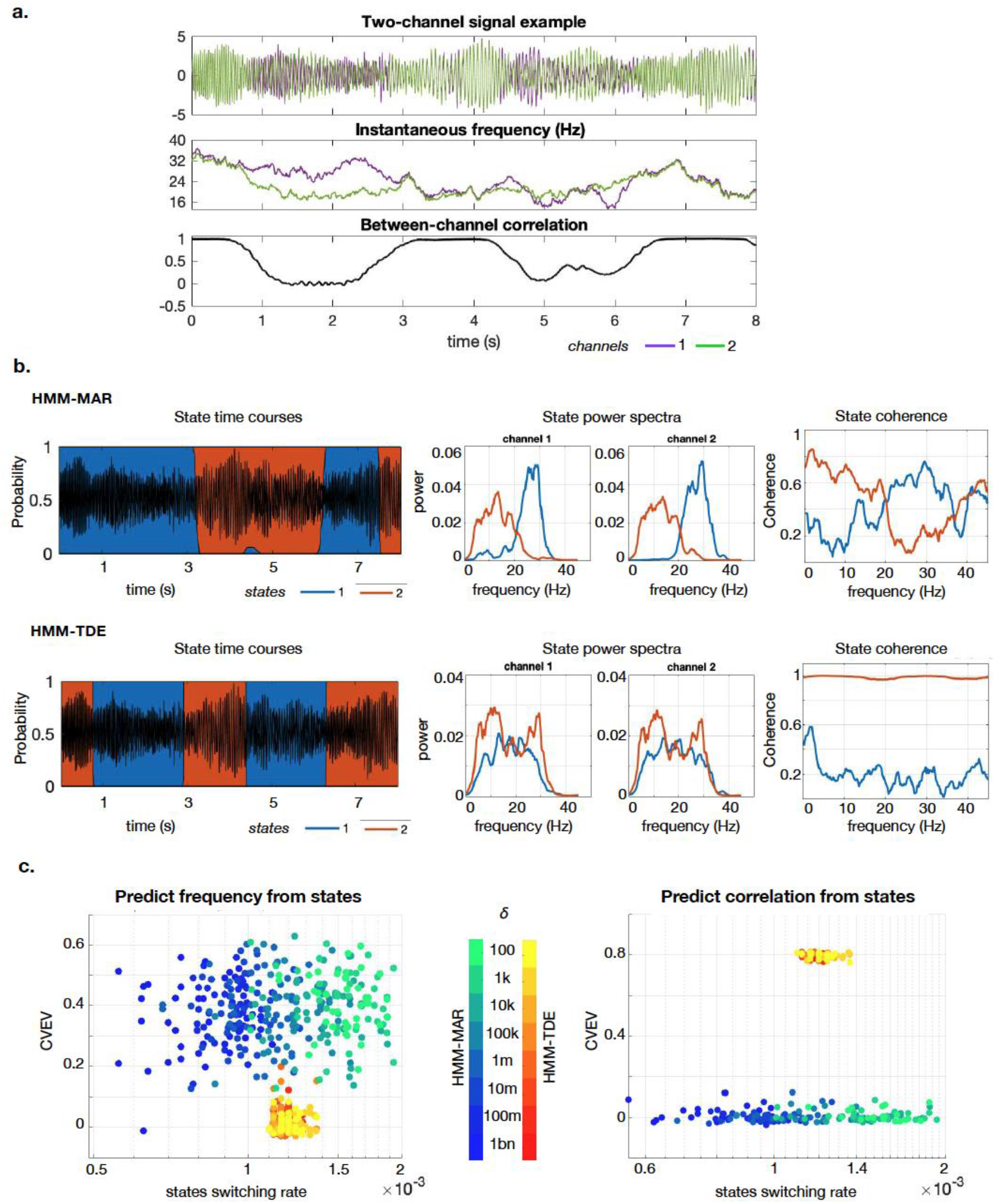
Experiments on two-channel synthetic data, with time-varying frequency, amplitude and broadband functional connectivity (expressed as between-channel correlation). **a**. Example of the synthetic signals; instantaneous frequency is shown in the middle panel, and instantaneous correlation in the bottom panel. **b**. On the left, examples of the state time courses of HMM-MAR (top panel) and HMM-TDE (bottom panel) applied to the signal in a; shown also the corresponding state power spectra (middle) and coherence (right). Here, *δ* = 10k, HMM-MAR order *P*=3, and the HMM-TDE lags *L*=15, with *S*=3. **c**. Cross validated explained variance (CVEV) of the HMM states predicting the ground truth instantaneous frequency of the channels (average explained variance across channels, left) and channel correlation (right) for 10 repetitions of the experiment and for different values of the state switching rate, manipulated through *δ*. Order and lags set as in **b**.

#### 2.1.2 Real data

##### LFP data

Wakeful resting state data were collected for 30 mins from a mouse’s hippocampus, using intracranial (Neuropixel) recordings. We used two channels from the array of 385 neuropixel electrodes for our tests. These data are part of a yet unpublished dataset, but are available upon request.

In detail, C57BL6/J male mice were first anesthetized with isofluorane (1% in oxygen) and placed in a stereotaxic apparatus (Kopf, California). A craniotomy was performed, centered in AP: -3.2, ML: 3 coordinates from bregma. Then, a head fixation crown (Neurotar, Helsinski) was implanted and secured with UV curating cement. Finally, a ground wire was implanted in the superficial layers of the cerebellum. The exposed craniotomy was covered with silicone (Kwik-cast, WPI) for protection. Meloxicam (5 mg/kg) were injected subcutaneously for three days after the surgery for pain and inflammation relief. The animals were allowed to recover for five days with food and water *ad libitum*. Then they were handled twice a day and placed in the head fixing apparatus for increasing times (10 mins to 40 mins), for 6 days to reduce stress. In the 7^th^ day, the animal was head-fixed, and the silicone removed from the craniotomy. Then, a Neuropixel 2b probe was inserted at 1μm/second. The probe was inserted a total of 4 mm corresponding to the first 385 recording sites of the probe. These spanned secondary visual cortex and ventral dentate gyrus. LFP data was filtered (0.5-1000 Hz), amplified and digitized (2.5 kHz). Data was acquired for 30 mins, while the animal was in resting state, in darkness. For our analyses, we used 2 channels, chosen such that their activity was not very correlated, from the hippocampus of one mouse and downsampled the data to 250 Hz.

All procedures involving animals were approved by the Danish Animal Experiment Inspectorate. Animals received food and water *ad libitum* and were housed under a 12h light-dark cycle. After surgery, animals were housed individually and allowed to recover for at least 3 weeks before experiments.

##### MEG data

The MEG data used in this study were collected by O’Neill *et al*. [23], where 8 subjects were instructed to perform a button press with the index finger of their left hand using a keypad, roughly every 30 seconds and without counting the time in between button presses. Total scanning time was 1200 seconds per subject. The data were acquired using a 275 channel CTF whole-head system (MISL, Conquitlam, Canada) at a sampling rate of 600 Hz with a 150 Hz low pass anti-aliasing filter. Synthetic third order gradiometer correction was applied to reduce external interference. The data were converted to SPM8 and downsampled to 200 Hz. We used the same preprocessing pipeline as in [21]. After the removal of artifacts related to eye-blink and heartbeat with Independent Component Analysis (ICA) [24], the data were band-pass filtered between 1 and 48 Hz, and source-reconstructed to the two primary motor cortices (M1).

The data acquisition was approved by the University of Nottingham Medical School Research Ethics Committee.

### 2.2 Models

#### 2.2.1 The Hidden Markov Model (HMM)

The Hidden Markov Model (HMM) is a family of probabilistic models describing time series data as a sequence of *K* states [10]. Each state corresponds to a different probability distribution (also known as observation model), belonging to a pre-specified family of probability distributions (e.g., Gaussian). The HMM inference estimates, in a data-driven fashion, the state parameters, the probability of each state being active per time point (*state time course*), the transition probability matrix (i.e., the probability of changing from one state to another, and of remaining in the same state), and the initial state probabilities (i.e., the probability of each state at trial start).

Here two types of HMM were explored, the HMM-MAR [21] and the HMM-TDE [12], each with a different observation model (see below). Both are implemented in the HMM-MAR toolbox, publicly available on GitHub^1^. In our analyses, we manipulated:

- The respective model hyperparameters: the order *P* for the HMM-MAR and the lags structure for the HMM-TDE, defined by the width *L* and the inter lags steps *S* (see below for definitions).
- The prior probability of remaining in the same state as opposed to moving to another state, parametrized by the Dirichlet distribution concentration parameter^2^, denoted as *δ*.
- The number of states *K*.

We used non-parametric estimations of the state spectra and coherence^3^.

#### 2.2.2 Observation Models

##### Multivariate Autoregressive (MAR) observation model

Given a multichannel time series ***y***, the MAR models the signal at each data point ***y***_***t***_ as a linear combination of previous time points (***y***_***t*−1**_, **…**, ***y***_***t*−*P***_), given the generative model

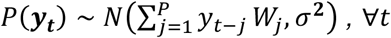

where ***W*** are the autoregressive coefficients, *σ*^*2*^ is the noise variance, and *P* is the autoregressive order, which determines the spectral resolution of the model. For a given choice of *P*, Bayesian inference is used here to estimate ***W*** and *σ*^*2*^, given the data.

In the HMM-MAR, each state’s probability distribution is represented by a set of coefficients ***W***^(***k***)^ per state *k*, as per:

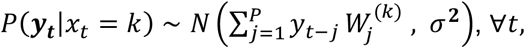

where *x*_*t*_ is a hidden variable indicating the state active at time point *t*; *x*_*t*_ is also estimated from the data.

When dealing with one-channel signals, we will just refer to the MAR model as autoregressive model or AR (and, correspondingly, HMM-AR).

##### Time-Delay Embedded (TDE) observation model

Instead of modelling the probability of observing every time point of the data, the TDE models the autocovariance of the signal around time point *t*. The relevant hyperparameter here is the lag structure, defined by *L* (the width of the signal’s window to consider at each time point) and *S* (how many steps separate each lag), as per *[-L, -L+S, …, 0, …, L-S, L]*. Mathematically, given the expansion ***Y***_***t***_ ***=*** (***y***_***t*−*L***_, **…**, ***y***_***t***_, **…**, ***y***_***t*+*L***_), the model is defined as Gaussian:

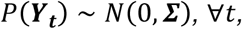

where ***∑*** is the autocovariance of the signal encoding linear relations across regions and time points within the window around *t*. The HMM-TDE is then defined as

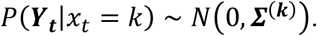

### 2.3 Analysis

Two types of analysis were performed: first, a sensitivity analysis on the observation models with respect to data features, as well as amount of training data and model hyperparameters; and second, a prediction analysis on the full HMM distributions assessing the generality of the estimation of the content of the states, as well as a permutation testing analysis to confirm the results.

#### 2.3.1 Sensitivity analysis on the observation models

The sensitivity of the two observation models, MAR and TDE, to different data features was tested: specifically, we assessed how the manipulation of frequency, amplitude and noise variance, as well as amount of data and model hyperparameters (the order for the MAR *P* and the TDE lags manipulated by *L*, while *S* was kept to 1) affected the estimations of ***W*** and ***∑*** respectively. Two independent sets of synthetic stationary data (as described in **Section 2.1.1**) were generated, both containing signals with all the above-mentioned feature combinations. One set was used for training the models and one for testing them. Since this analysis involved one-channel stationary signals, we will refer to the MAR model as AR. For every signal ***y*** from the training set, the model coefficients ***W*** (for the AR) and ***∑*** (for the TDE) were estimated. We will refer to these as ***W***(***y***), and ***∑***(***y***). The trained models were then tested on one target signal ***z*** from the test set. The capacity of the models to describe this test signal was measured by means of the log-likelihood ratio (of trained vs tested signal): this gave an indication of how similar training and test signals were, according to the trained models. Being model-dependent, the log-likelihood ratio expresses the sensitivity of each model with respect to each feature.

For the AR model, the likelihood of the target test signal ***z*** to be distributed as a training signal ***y*** is:

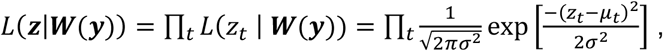

where 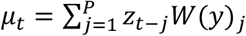 is the prediction from the autoregressive model, and *σ* is the noise standard deviation, also estimated from the data.

For the TDE model, given ***Z***_***t***_ ***=*** (***z***_***t*−*L***_, **…**, ***z***_***t***_, **…**, ***z***_***t*+*L***_), the likelihood of ***Z***_***t***_ to be distributed according to *N*(0, ***∑***(***y***)), for a training signal ***y***, is:

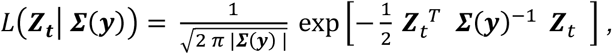

where |***∑***(***y***)| indicates the determinant of ***∑***(***y***). Then, the likelihood across time points becomes:

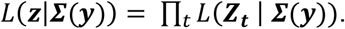

Finally, considering the likelihood of target signal ***z*** to be parametrized by a model trained on ***z*** (i.e., a training signal with the same features), *L*(***z*|*W***(***z***)) and *L*(***z*|*∑***(***z***)), the log-likelihood ratios are defined as:

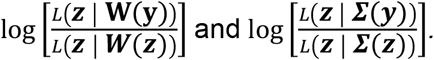

The log likelihood ratio is therefore a measure of the precision of the model in describing the test signal, given the training signal. If the log likelihood ratio of the model trained on ***y*** vs ***z*** is low (i.e. a large negative number), then the parameters of the model fitted to ***y*** do not describe well the test data ***z***, and, consequently, the two signals are regarded as different according to the assumptions of the model; on the other hand, if the log likelihood ratio is close to zero, both the models trained on ***y*** and ***z*** describe well the test data and, therefore, that the two signals cannot be distinguished given the model definition and assumptions.

#### 2.3.2 HMM analyses on synthetic data

The previous analyses were designed to compare the observation models as stand-alone distributions. Next, these were assessed within the HMM framework. Specifically, the HMM states were analyzed by regressing the state time courses on the ground-truth data features — instantaneous frequency *f(t)*, amplitude *a(t)* and between-channel correlation *r(t*). In order to use a cross validation scheme, the data, simulated as a unique session of *T*=50000 points, were partitioned into trials of equal length. The cross validation was hence performed at the trial level, grouping the trials into 10 folds (using 9 for training and the remaining 1 for testing, in turns). This yielded a measure of accuracy —cross-validated explained variance (CVEV)— per feature, describing how well the HMM states captured that feature. The accuracy of the models in capturing each feature was tested as a function of the HMM hyperparameters (order for the HMM-MAR and lags for the HMM-TDE), number of states, and of the prior probability of remaining in the same state, modulated by the hyperparameter *δ* (see above). To confirm the reproducibility of the results, each analysis was repeated 10 times.

A permutation testing analysis was conducted (10000 permutations) to further validate the effects of the prediction analyses, where the null hypotheses are of the sort of “the HMM states do not capture amplitude”. The permutations were also performed across trials.

#### 2.3.3 HMM analyses on real data

The two HMMs were also tested on LFP and MEG real data, comparing the frequency properties of the states. For the MEG data, the state temporal information was also contrasted to the available task data.

### 2.4 Code accessibility

All the code used for the generation of synthetic data and for the analysis of both synthetic and real data will be available upon publication on Github^4^.

## 3. RESULTS

We first describe the behavior of the TDE and AR as standalone models with respect to different data characteristics, and then we describe them in the context of the HMM.

### 3.1 Observation models

In the first part of this study, a sensitivity analysis was conducted on the two HMM observation models (AR and TDE), with respect to data characteristics and model hyperparameters, which we first considered as standalone models. The models were trained and tested on synthetic, stationary sinusoids (described in **Section 2.1.1**, see examples in **Figure 2a**), and their sensitivity measured by the log likelihood ratio of train vs. test signal, which we can interpret as a measure of how well the models could identify signal differences (see **Section 2.3.1**).

To test the model sensitivity to frequency, the models were trained on signals that only differed in frequency (i.e., with a fixed amplitude and noise variance). We then selected a given target test frequency and tested the models (trained for multiple frequencies) on the test signal, which had the same amplitude and noise variance as the training signals. The procedure was repeated to test different amplitude and noise values (**Figure 2b**). The TDE model showed lower frequency resolution than the AR model (note the different x-axes scales in the plots). Generally, decreasing amplitude and increasing noise had a similar effect on the sensitivity of the models to frequency. For the AR model, signal-to-noise ratio (SNR), rather than amplitude or noise variance as separate factors, affected its frequency resolution: that is, the smallest detectable frequency difference depended on the signals’ SNR. This intuition can be verified analytically by proving that the coefficients of two models trained on signals with different amplitude and noise variance, but same SNR value, are the same (see **Appendix**).

To test the model sensitivity to amplitude, the same procedure detailed above was adopted: the models were trained and tested on signals that only differed in amplitude, for some fixed values of frequency and noise variance (**Figure 2c**). While the AR model clearly distinguished the full range of amplitudes tested, it was more difficult for the TDE to distinguish training and test signal, especially when the training amplitude was higher than the test one. Because of this, we tested a wider range of amplitude differences for the TDE model, where we used multipliers of the target test amplitude as training amplitudes (for the AR model, the different training amplitudes were linearly spaced). Overall, amplitude resolution was not affected by frequency, as the model relative performance was similar for all tested frequencies.

Frequency and amplitude resolution of the models were further investigated by varying the amount of training data (manipulating signal length) and the model hyperparameters —see **Supplementary Figure 1**. Briefly, more training data induced a higher resolution on frequency in both models, and a higher sensitivity to amplitude in the AR model. Also, a larger lag window in the TDE model increased frequency resolution. Neither a higher amount of data, nor a bigger lag window significantly changed the TDE sensitivity to amplitude. As for the AR model, since an autoregressive order of 3 can already capture one fundamental frequency [22], and because in this particular case the data only had one frequency, an increase in the autoregressive order did not change the model performance. That is, while a higher autoregressive order could explain more complex, multi-frequency spectral patterns, it does not improve the resolution of a single frequency.

In summary, the AR model accurately described stationary sinusoids with sufficiently high SNR and showed high frequency resolution, which was also influenced by the amount of training data. The AR model was comparably less sensitive to amplitude than to frequency, and also amplitude resolution was influenced by the amount of training data. On the other hand, the TDE model showed a lower frequency resolution than the AR. This resolution was modulated by the lag hyperparameter and by the amount of training data. Finally, the TDE model was less sensitive to differences in amplitude, and, unlike the AR, its sensitivity was not symmetric (see **Figure 2c**); this is because, for the TDE (i.e., for the Gaussian distribution), the log-likelihood function (unlike the probability density function) is not symmetric with respect to the scale. Manipulations in lags or amount of training data did not significantly change this behavior.

### 3.2 HMM inference

Next, the HMM-MAR and HMM-TDE capacity to represent dynamic changes in the signal was investigated, using synthetic, non-stationary signals with time-varying frequency, amplitude and functional connectivity (as described in **Section 2.1.1)**.

#### 3.2.1 Detection of changes in frequency and amplitude

The HMM-MAR and the HMM-TDE were tested on single-channel signals with time-varying frequency and amplitude (first with single frequency, then with two concurring frequencies). As mentioned in **Section 2.1.1**, the generative model of the signals, unlike the HMM’s, does not assume quasi-stationary periods of sustained oscillations, and is, therefore, more general. While we could have generated data that followed the assumptions of the HMM, real electrophysiological data is likely to not follow these assumptions, so we opted for a more assumption-free generative model. In detail, one-channel, non-stationary signals were fed to the HMMs and the resulting state time courses (i.e., the probability of each state being active at each time point) were subsequently analyzed. In particular, standard regression was used to predict the frequency and amplitude time courses from the state time courses, in a cross validated fashion. The prediction analysis was performed varying the hyperparameter *δ* (related to the prior probability of remaining in the same state, which influences the state switching rate; see above), as well as the number of states *K*, the order *P* for the HMM-MAR and the lags (*L* and *S*) for the HMM-TDE. The procedure was performed 10 times per configuration (see **Section 2.3.2**).

When the signals varied in frequency and amplitude with one single frequency active at any given time (see example signal in **Figure 3a**), both HMM-MAR and HMM-TDE clearly captured frequency changes, with each of their states representing a different frequency band (**Figure 3b**). Our prediction analysis showed that, for fixed order and lags, the state switching rate affects model performance in capturing frequency (see **Figure 3c**, where order *P*=3, lags structure set to *L*=15, spaced in steps of *S*=3): clearly, the models with faster switching rates (here, the HMM-TDE runs) could explain frequency variance better (up to 90%), meaning that they better matched the dynamics of the data. For comparison, a separate permutation analysis showed that randomly assigned states could explain only 0.0132, ± 0.0130 of frequency variance (CI = 0.95, p < 0.0001 for all models). The prediction analysis further showed that varying the model hyperparameters also had an influence on the performance of the models and on the state switching rate. More specifically, while *δ* had a great influence on the HMM-MAR performance, widely modulating the state switching rate, the HMM-TDE performance and state switching rate were more affected by the lags manipulation than by *δ* (see **Supplementary Figure 2a** for further details). None of the models could explain amplitude variance significantly better than randomly assigned states. To assess whether these conclusions were contingent on the tested frequency, the models were run on signals spanning two different frequencies (alpha and gamma), one frequency band at a time. Our results hold similarly for both frequency bands; see **Supplementary Figure 3**.

To further investigate the extent to which the HMM states could capture amplitude, the same analysis was performed on one-channel, non-stationary signals only varying in amplitude (a signal example is reported in **Figure 3d**). This time, coarse changes in amplitude could be captured using two states for each model (**Figure 3e**). The prediction analysis (now regressing the state time courses on the amplitude time course) for a fixed order and lag choices (as before, *P*=3, *L*=15, *S*=3), showed that both models were able to detect amplitude changes in this scenario, but the extent to which they could do so depended on how well the state switching rate matched the ground-truth dynamics of the data (**Figure 3f**); for example, the extreme cases when the switching rate was close to zero (leftmost in the panel) corresponded to cases when one state dominated the entire decomposition, driving the CVEV to zero. This is in contrast with the previous analyses, where we examined the states as stand-alone distributions, and where the AR model was considerably better at capturing amplitude. The difference here is however at the level of the HMM inference, with certain HMM-MAR runs not being able to switch states at sufficient speed (or even collapsing to a single state) for some choices of *δ*. For comparison, randomly assigned states explained 0.0228 ± 0.0227 (CI = 0.95) of the signals’ amplitude, yielding statistical significance for all decompositions that did not degenerate onto a single state. The complete analysis results can be found in **Supplementary Figure 2b**.

We also explored the HMM-MAR and HMM-TDE inference for a higher number of states, using signals that vary both in frequency and in amplitude (**Supplementary Figure 4**). When increasing the number of states, the HMM-MAR started capturing amplitude instead of only frequency (**Supplementary Figure 4c**). In contrast, endowing the HMM-TDE with more states resulted in a higher frequency band resolution, but no better sensitivity to amplitude (**Supplementary Figure 4f**).

We also tested the models on synthetic data with two concurring oscillations: one with higher power in low frequency, and another with lower power in high frequency (see **Figure 4a** for an example, and **Section 2.1.1** for data generation details). The power ratio between the two frequency bands was varied from 4 (i.e., the low frequency has four times the power of the high frequency) to 1 (i.e., equal power). For each power ratio, we generated data and ran the HMM-MAR and the HMM-TDE ten times for each choice of the number of states (*K*=2,3,5). Then, the signals’ frequency and amplitude in both frequency bands were predicted from the states time courses in a cross validated fashion. So, for each HMM run, four decoding analyses were made, predicting: (i) frequency and (ii) amplitude in the low frequency band, and (iii) frequency and (iv) amplitude in the high frequency band (see **Figure 4b and c** for HMM-MAR and HMM-TDE states time courses and power spectrum example). As shown, different power ratio between the frequency bands changed the data and, accordingly, the HMM behavior. When the power ratio between low and high frequency band was higher, HMM-TDE states captured mostly frequency in the low frequency band (see **Figure 4d**, leftmost panel, bottom); however, when a more equal distribution of power between bands was used, the HMM-TDE states, given its configuration of lags, were better at capturing frequency changes in the higher frequency band (see **Figure 4d**, progression of bottom panels). On the other hand, the HMM-MAR states exhibited a more mixed description of the signals, capturing both frequency and amplitude in the two frequency bands even for the highest power ratio (see **Figure 4d**, leftmost panel, top). This changed toward a preference of frequency in the higher frequency band when the power ratio approached 1 (see **Figure 4d**, rightmost panel, top). Note that these conclusions depend to some extent on the parametrization of the MAR and TDE models; for example, if we had more spaced lags, the HMM-TDE sensitivity would lean more to slower frequencies. The choice of K, however, did not affect these conclusions.

In conclusion, when using signals with one single frequency component (with dynamically varying frequency and amplitude), both models were more sensitive to frequency than they were to amplitude (independently of the frequency band of the signal), but they could also capture changes in amplitude when frequency was relatively constant and, in case of the HMM-MAR, when endowed with a higher number of states. How well the decomposition captured differences in amplitude and frequency depended on the temporal dynamics of the state time courses (i.e., the state switching rate) and how well they matched the underlying modulations in the data. Although the main conclusions stand with more than one dynamically changing frequency component, the HMM-MAR exhibited in this case a more complex behavior. As shown, the switching rate can be manipulated by modifying the observation model hyperparameters (*L* and *P*) and the priors of the transition probability matrix (*δ*). In real scenarios, these hyperparameters could be tuned to access different temporal scales in the data.

These analyses may seem opposed to the previous section, where the MAR model showed to have a higher resolution in both frequency and amplitude. These findings can be reconciled by the fact that the MAR has a higher capacity to explain variance in the raw data, and thus, when plugged into the HMM inference, might necessitate less state switching (specially, when overparametrized). That is, while a high-order single autoregressive model could potentially explain the data very well and therefore dominate the decomposition, the corresponding state time courses would not be effective to describe the time-varying facets of the data, which is what our CVEV metric precisely captures. While the same problem could occur for the HMM-TDE as well, it is less likely to happen because the number of effective parameters of the TDE (and therefore its sensitivity to the different features, as shown in the previous section) is lower.

#### 3.2.2 Detection of changes in functional connectivity

Next, The HMM-MAR and the HMM-TDE were tested on synthetic two-channel, non-stationary signals with time-varying correlation across the entire frequency spectrum (see **Section 2.1.1**), which we will refer to as *broadband functional connectivity* (see **Figure 5a** for examples of signal and between-channel correlation). Again, a regression analysis was used to predict frequency and between-channel correlation from the state time courses, varying *δ* and the observation model hyperparameters. The full experiment was repeated 10 times.

Here, the HMM-MAR primarily captured frequency information specific to the single channels, whereas the HMM-TDE was able to capture broadband functional connectivity, such that one state was assigned to the periods of highest correlation between the two channels and the other state to periods of lower correlation. This can be qualitatively observed in the state time courses (**Figure 5b**, leftmost panels) with respect to the channel correlations (**Figure 5a**, bottom panel), and in the estimation of the state power spectra (**Figure 5b**, middle panels) and the state coherence (**Figure 5b**, rightmost panels). The prediction analysis quantitatively corroborated this, showing that, for fixed order and lags (here, *P*=3 and *L*=15, in steps of *S*=3), varying *δ* to manipulate the state switching rate did not change the HMM-TDE ability to capture broadband functional connectivity significantly (**Figure 5c**, right panel). This analysis also revealed that the HMM-MAR performance in predicting channel frequency was not very stable across runs (see how scattered the performance is along the y axis in **Figure 5c** left, regardless of the *δ* parameter). For comparison, a permutation testing analysis resulted in randomly assigned states explaining 0.0126 ± 0.0125 (CI=0.95) of the signal frequency variance, and 0.0029 ± 0.0027 (CI = 0.95) of channel correlation variance; this means that the HMM-TDE did not perform significantly better than randomly assigned states in capturing frequency, and neither did the HMM-MAR in capturing broadband coherence.

We repeated the prediction analysis by varying the model hyperparameters (complete results in **Supplementary Figure 5**), showing again that the HMM-MAR performance and state dynamics were more affected by changes in *δ* than HMM-TDE, which was most affected by the lag configuration.

But how can these results be reconciled with previous work where the HMM-TDE was successfully used to find states with distinct amounts of coherence in different frequency bands (for example, in [12, 25, 26])? First, our analyses only used two states, inducing the models to focus on what was most salient to them. Note that while we could have used more states, the goal of this paper was precisely to characterize the models’ saliency to the different data features. Since the HMM-TDE has less spectral detail than the HMM-MAR, broadband coherence resulted to be its most salient feature. Second, power and coherence are not independent. The HMM-MAR’s states had different spectral signatures in coherence (**Figure 5c**, right-bottom panel) also because they reflected their respective spectral signatures in power (**Figure 5c**, middle-bottom panel). We established before that the HMM-MAR has great sensitivity to frequency changes, which were present in this data (**Figure 5a**, middle panel) and hence captured by the model. Altogether, our results do not imply that the HMM-TDE is unable to find spectral-specific changes in coherence, if they existed in the data, but that it would do so with less spectral detail than the HMM-MAR and it would instead prioritize broader frequency bands (or broadband coherence as in this example).

In summary, for bivariate signals exhibiting intermittent periods of high correlation, the first feature of focus for HMM-TDE was broadband connectivity, while the HMM-MAR was more sensitive to detailed frequency modulations and less to broadband connectivity.

#### 3.2.3 Real data

We ran the two HMM on two real datasets: LFP data from mice in wakeful rest, and MEG data of humans performing a simple motor task (source-reconstructed onto two motor cortex parcels); see **Section 2.1.2** for details.

**Figure 6a** shows the spectral properties of the LFP data. Here, the state time courses of HMM-MAR and HMM-TDE were relatively well correlated, with an average correlation of 0.5, for the various tested configurations (number of states *K*=2,3,4, HMM-MAR order *P*=3,5,7,9; HMM-TDE lags *L*=3,5,15,21 in steps of 1, *δ* = 1k, 10k, 100k, 1m, 10m), indicating that the decompositions had overlapping properties. However, we found that the HMM-TDE inference exhibited much less uncertainty, namely that the HMM-TDE inference assigned very high probabilities to one state at each time point, while the HMM-MAR estimates had a larger uncertainty (with the state probabilities more rarely reaching values close to 1.0); see state time courses and state properties of HMM-MAR and HMM-TDE in **Figure 6bc**. In particular, the HMM-TDE assigned probability of ∼1.0 to 90% of the time points on average, while for the HMM-MAR less than 20% of the points had a state with probability near to 1.0. This uncertainty likely follows from the rich frequency content of high-quality LFP combined with the higher spectral sensitivity of the HMM-MAR.

**Figure 6.**
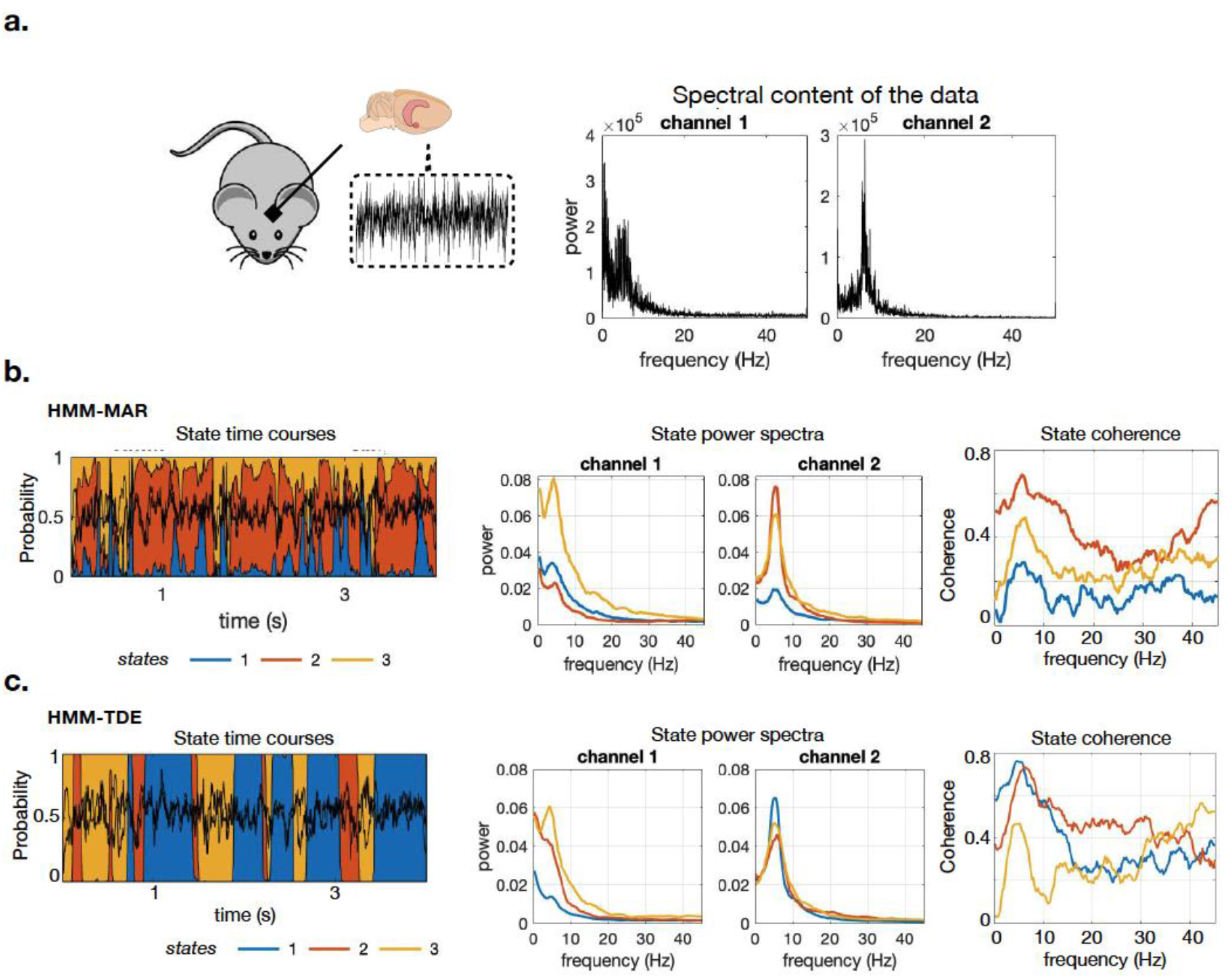
Experiments on LFP data, with two LFP channels (chosen such that their activity was not very correlated) from the hippocampus of a mouse during resting state, downsampled to 250 Hz. **a**. Spectral content of the two channels. **b**. Example of HMM-MAR state time courses, as well as the state power spectra and the state coherence. **c**. Similar to **b**, for HMM-TDE. The models are trained on 30 mins of data with three states (here, HMM-MAR order *P*=5, HMM-TDE, lags *L*=15, *S*=3, *δ*=100k).

When applied to MEG data (see **Figure 7a** for a spectral characterization of the data), Vidaurre *et al*. [21] previously showed that the HMM-MAR could capture task-related information without the model having prior knowledge of the task. Here, we ran the HMMs subject by subject and, similarly to [21], computed the response *evoked state probability* (i.e., the average probability of each state to be active within a window around the finger tapping event), alongside with the state power spectra. Since the channels were made orthogonal in order to correct for signal leakage [27] their coherence was greatly diminished (and not considered here). Like with the LFP data, the two HMM variants differed in the uncertainty of the state assignments, with the HMM-TDE assigning probabilities more sharply than the HMM-MAR (see **Figure 7b** and **c**, leftmost panels, for an example of the HMM-MAR and HMM-TDE state time courses). Still, both the HMM-MAR and HMM-TDE captured task information well; specifically, one state showed increasing probability of being active in the 2 seconds before the button press and has a drastic drop 1-2 seconds after, while another state had a specular behavior, with very low probability of being active before the button press and a sharp increase 2-4 seconds after (see **Figure 7b** and **c**, rightmost panels, with the response-evoked state probabilities for HMM-MAR and HMM-TDE respectively). **Figure 7** shows the results for one exemplary subject; see **Supplementary Figure 6** for the other subjects.

**Figure 7.**
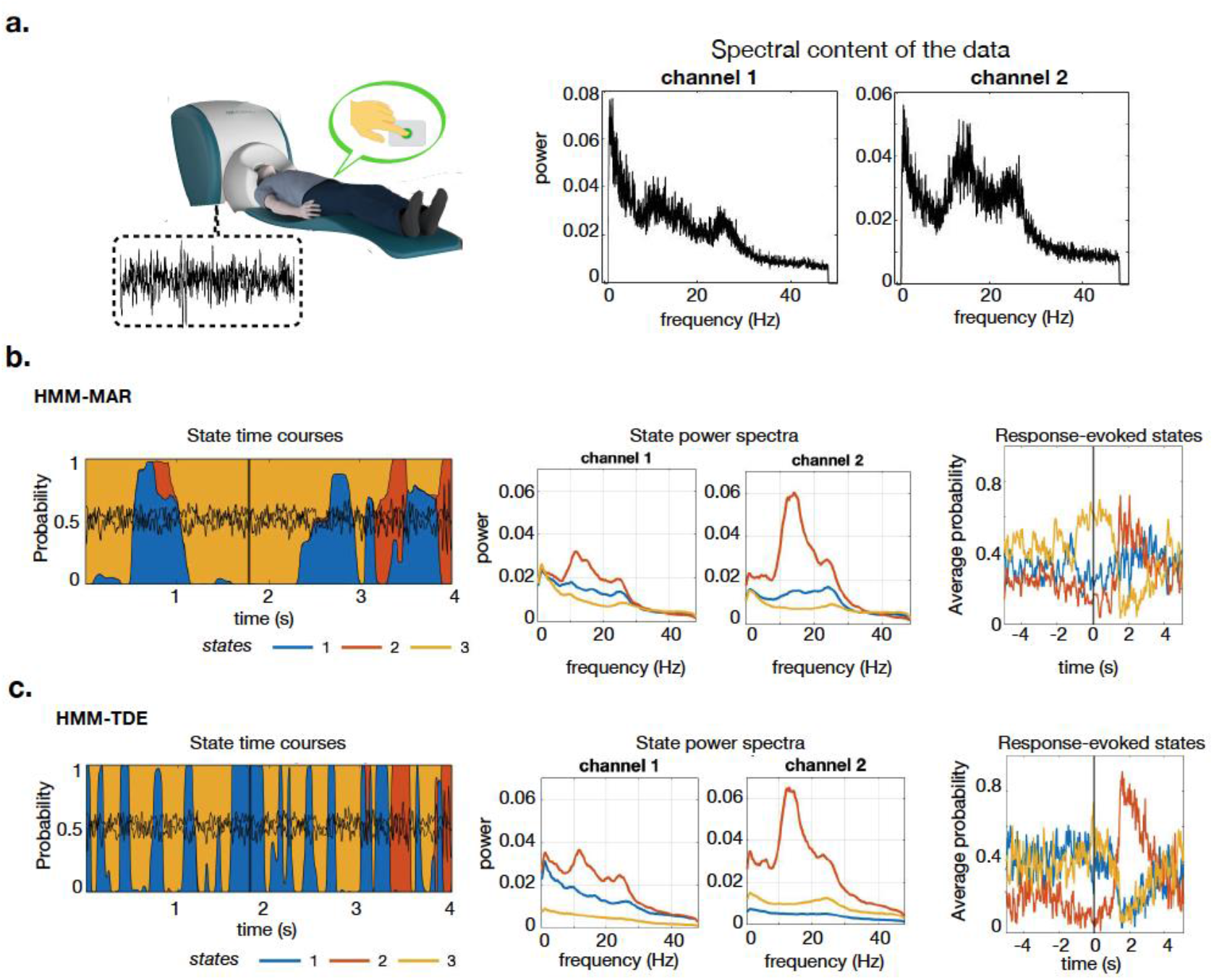
HMM-MAR and HMM-TDE applied to MEG data. **a**. The data for this analysis are 2 MEG channels from the motor cortex of one (human) subject who performed a simple finger tapping task. Data were downsampled to 200 Hz and band-filtered between 1 and 48 Hz. The spectral content of the two channels is also shown. **b**. Example of HMM-MAR state time courses, around a task-relevant moment (button press), marked by a black vertical line (left); the corresponding states power spectra (middle); and the probability of states around the button press (response-evoked states probability, rightmost panel). **c**. Same as in **b**, for the HMM-TDE model. The models are trained on 20 mins of recordings with three states. For the HMM-MAR we used order *P*=3, and for the HMM-TDE we used *L*=1, with *S*=1. *δ*=100k.

In conclusion, here the state spectral properties were not as different between two models as in the synthetic simulations, but the HMM-TDE performed in a more decisive way in the sense of having a higher certainty in the state time courses estimation (see **Supplemental Results** for further analyses on how the dynamics of the signal affects uncertainty of the models).

## 4. DISCUSSION

In this study, focusing on electrophysiological data with one or two channels —both real and simulated, we explored in detail the behavior of two types of HMMs: the HMM-MAR [21] and the HMM-TDE [12]. We excluded other models that are instead run on amplitude time series derived from, for instance, a Fourier transform [14].

We did not consider high-dimensional data (i.e., with many channels), where the number of parameters of the HMM-MAR scales quadratically on the number of channels, so it soon becomes overparametrized; in this case, HMM-TDE is preferred, as discussed in [12]. Using synthetic data, we first measured the sensitivity of the two observation models to different data features in a stand-alone manner. Then, we studied what dynamic aspects of the data drive the inference of the HMMs. Finally, we confronted the models with real MEG and LFP data. We note that this paper was focused on the signal properties of the data, so we were agnostic to which experimental paradigms (e.g. task or rest), behavioral states (wakefulness or sleep) or processing steps (whether band-pass filtering was applied) could give rise to specific types of modulations in the data.

Our sensitivity analysis of the standalone observation models showed that the MAR is generally more sensitive to frequency than it is to amplitude, and that it has a higher frequency resolution than the TDE observation model. We also found that it is harder for the TDE model to distinguish amplitude, and that its amplitude resolution is not symmetric (that is, the TDE is more sensitive to a testing amplitude that is higher than the training amplitude, and less sensitive to a testing amplitude that is lower than the training amplitude). The HMM analyses on single-channel signals with non-stationary properties and one single frequency active at a time showed that both the HMM-MAR and HMM-TDE preferentially capture changes in frequency when the signal varied both in frequency and in amplitude. Except for degenerate solutions, the HMM-MAR could capture changes in amplitude when the signal did not vary much in frequency, or when the model was endowed with a high number of states. The HMM-TDE successfully captured amplitude reliably when the signal was only varying in amplitude, provided that the changes in amplitude were large enough. HMM-TDE showed generally more robust states dynamics than the HMM-MAR. Finally, we found that their performance depended on hyperparameter configuration (on both the prior probability of remaining in the same state and on the model-specific hyperparameters). When two concurring frequency oscillations were present, the HMM-MAR’s behavior was more mixed, capturing not only frequency but also amplitude to some extent, while the HMM-TDE focused mostly on frequency. It is important to point out that the HMM-TDE behavior might change to some extent if PCA was applied beforehand, biasing it to higher frequencies when these are within the included principal components [28]. On two-channel synthetic data with transient periods of high functional connectivity (in the sense of between-channel correlation), we found that the HMM-MAR focuses first on representing frequency-specific changes at the expense of broadband functional connectivity (here, correlation), while the HMM-TDE clearly prefers focusing on broadband changes in functional connectivity above and beyond fine-grained fluctuations in the frequency content of the signals. Overall, we can conclude from these results that the HMM-MAR is the most appropriate model when the main goal of the analysis is detecting detailed changes in the frequency of the signal, or amplitude if the frequency is relatively stationary (for example as a result of filtering), while the HMM-TDE may be more appropriate in multi-channel data when the focus is on capturing spectrally-wide changes in functional connectivity, and, in general, when the MAR model is overparametrized and the inference collapses onto one single state.

Having a greater sensitivity to the different features present in the data may however come at a price. We here referred to *uncertainty* of the HMM inference as how decisive was the state assignment: i.e., if state probabilities are 1/K for all states at time point t, that would be the most uncertain; whereas if one state has probability 1.0 and the rest have 0.0, then this would be the most certain assignment. Although potentially related, this is a different measure to the variability in the estimation that the states time courses might exhibit over multiple runs (see [29]). Whereas the HMM-MAR is more sensitive to detailed spectral content than the HMM-TDE, this can also result in greater estimation uncertainty, or even degenerate solutions when only one state is active. This has made the HMM-TDE the model of choice in several applications of the HMM in electrophysiological data [30, 31, 18, 26]. Indeed, uncertainty can translate in lower reproducibility of the results, since small changes in the data are more likely to elicit larger changes in the resulting estimates. The cause of HMM-MAR’s greater uncertainty is likely due to the nonlinearity of neural oscillatory signals, in the sense that the phase of the oscillations does not progress at linear steps, and, therefore, the shape of the wave is often irregular and asymmetric (that is, the instantaneous frequency of the signal varies irregularly from time point to time point [32]). Since both the TDE and the MAR models are linear, the only way to accurately model abrupt changes in instantaneous frequency is through the state time courses. For this reason, over-parametrized models with excessive sensitivity can be more volatile, and can even exhibit quick state switches within single oscillatory phase. Whereas these estimates are technically not incorrect, they lend to more complex interpretations.

While the balance in sensitivity to frequency and amplitude for the different models was unambiguously characterized in our single-channel experiments, it is important to note that our analyses are to some extent contingent to the generative model used to sample the data and its assumptions. In particular, our generative model (as described in **Section 2.1.1**) for pairs of channels did not explicitly model spectrally fine-grained changes in coherence, and instead manipulated spectrally broadband changes. Here, the HMM-TDE was clearly focusing mostly on changes in functional connectivity, but the HMM-MAR would have probably been more sensitive to connectivity if the ground-truth generative model were targeted at producing frequency-specific changes in coherence. While our generative model for sampling data is a reasonable approximation of actual empirical data, the sensitivity of the two models to connectivity will in the end be determined by how much explained variance in the real data is attributable to changes in coherence vs. changes in power, and how this variance is spectrally distributed (i.e., coarse vs. fine-grained). We also note that the procedure we used to simulate the data was driven by practical reasons, such that we could better quantify which aspects of the signal are preferentially captured by the HMM. Some real data have features, like their 1/f nature, that are not reflected in our simulated data. Our conclusions are however focused on the models’ sensitivity, not on the specifics of the data, and therefore hold generally.

Finally, one frequently asked question about HMM analyses is how to choose the number of states. Our conclusions were not critically influenced by this. In general, if the goal is interpretation, parsimony is a reasonable criterion: if fewer states are sufficient to capture the features of interest, this might be preferrable. On the other hand, if a finer description of the data is needed, then the number of states may be increased. An example of this was shown in **Supplementary Figure 4**, where a higher number of states was needed to detect changes in amplitude above and beyond changes in frequency.

## 5. CONCLUSION

Unsupervised methods of analysis provide a useful tool for discovery and are freer of researcher bias than other methods. But, at the same time, they can be a black box in the sense that we do not precisely know what aspects of the data they capture. Here we focused on an unsupervised method often applied to electrophysiological data, the hidden Markov model. Our aim was to characterize precisely what aspects in the data, such as frequency or amplitude modulations and functional connectivity, drive an HMM estimation. Using synthetic as well as real (LFP and MEG) data, we characterized the behavior of two different types of HMM (the HMM-MAR and the HMM-TDE). In summary, we showed that both HMMs preferentially capture frequency modulations, but the HMM-MAR does it in more detail —this, in turn, results in more decisive estimations for the HMM-TDE. For the same reason, the HMM-TDE is more effective in capturing functional connectivity modulations in relatively broad frequency bands. We note that the HMM is not a biophysical model, and different parametrizations or model choices offer just alternative perspectives of the data. On these grounds, none of these can be said to be biologically more or less valid than the others —only more or less practically useful given the characteristics of the data and the research goal.

## ACKNOWLEDGEMENTS

DV is supported by a Novo Nordisk Foundation Emerging Investigator Fellowship (NNF19OC-0054895) and an ERC Starting Grant (ERC-StG-2019-850404). MWW’s research is supported by the [NIHR](http://dx.doi.org/10.13039/100006662) Oxford Health Biomedical Research Center, the Wellcome Trust (106183/Z/14/Z, 215573/Z/19/Z), the New Therapeutics in Alzheimer’s Diseases (NTAD) study supported by UK MRC, the Dementia Platform UK (RG94383/RG89702) and the EU-project euSNN (MSCA-ITN H2020–860563). This research was funded in part by the Wellcome Trust (215573/Z/19/Z). For the purpose of Open Access, the author has applied a CC BY public copyright licence to any Author Accepted Manuscript version arising from this submission.

## Conflict of interest statement

All authors declare that they have no conflicts of interest.

## Footnotes

1 https://github.com/OHBA-analysis/HMM-MAR

2 In the HMM-MAR toolbox, this is specified with the *DirichletDiag* option.

3 Function *hmmspectramt()* in the toolbox.

4 https://github.com/LauraMasaracchia/HMM_explore

## Notes

### Competing Interest Statement

The authors have declared no competing interest.

### Summary of Updates

We have performed a new analysis on simulated data that represent electrophysiological data more realistically (new Figure 4). We have also performed a new analysis on the uncertainty of the HMM inference, reported in Supplementary Figure 7 and discussed in Supplemental Results. We have performed a new analysis on the frequency sensitivity of the HMMs per frequency band, reported in Supplementary Figure 3. Finally, we have elaborated on the choice of the states number K.

