## Appendix and supplementary figures for "Dissecting unsupervised learning through hidden Markov modelling in electrophysiological data"

We show that the (M)AR model is sensitive to SNR by proving that the coefficients of an autoregressive model describing a signal $\boldsymbol{y}$, with amplitude $a_y$ and Gaussian noise $\boldsymbol{\xi}_y$ with noise variance $v_y$, are the same of those describing a different signal $\boldsymbol{z}$, with different amplitude $a_z$ and different noise $\boldsymbol{\xi}_z$ with noise variance $v_z$, provided that the two signals have the same SNR.

Let us define the two sinusoidal signals as:

$$\boldsymbol{y} = a_y \sin(2\pi f \boldsymbol{t}) + \boldsymbol{\xi}_y,\ \boldsymbol{z} = a_z \sin(2\pi f \boldsymbol{t}) + \boldsymbol{\xi}_z,$$

where **t** = (1, …, T) is a row vector of length T. We set $a_z = 100\ a_y$, $v_z = 100\ v_y$, and hence $\boldsymbol{z} = 100\ \boldsymbol{y}$, and $SNR_y = SNR_z$.

Given the order *P* of the autoregressive models, we define **Y** and **Z** as the sets of *P* points preceding every time point *t* in $\boldsymbol{y}$ and $\boldsymbol{z}$ respectively. So **Y** and **Z** are matrices of dimension Tx*P*, where each row is:

$$\boldsymbol{Y_t} = (y_{t-P}, \ldots y_{t-1}),$$
$$\boldsymbol{Z_t} = (z_{t-P}, \ldots z_{t-1}) = 100\ \boldsymbol{Y_t}\ .$$

We finally find the autoregressive coefficients that predict $\boldsymbol{y}$ and $\boldsymbol{z}$ :

$$\boldsymbol{W_y} = (\boldsymbol{Y^T Y})^{-1}\boldsymbol{Y^T y} \quad \text{and} \quad \boldsymbol{W_z} = (\boldsymbol{Z^T Z})^{-1}\boldsymbol{Z^T z} = \frac{1}{10000}(\boldsymbol{Y^T Y})^{-1} 10000\ (\boldsymbol{Y^T Y}) = \boldsymbol{W_y},$$

which trivially proves that the autoregressive coefficients of two signals with different amplitude, but same SNR, are the same.

### SUPPLEMENTAL RESULTS

To investigate how the HMM uncertainty related to the signal dynamics, the HMMs were run on synthetic signals with slow-changing dynamics (spanning a frequency range between 0.1 and 45 Hz, and an amplitude range between 0.1 and 10). Uncertainty was measured as the proportion of timepoints with a state probability lower than 0.9. The final uncertainty measure was obtained averaging across 10 repetitions of the analysis. Each analysis was performed for three different values of $\delta$. As shown in **Supplementary Figure 7**, when applied to a signal with slow-changing dynamics, the uncertainty of the HMM-MAR estimates decreased to a level similar to the HMMTDE.

### SUPPLEMENTARY FIGURES

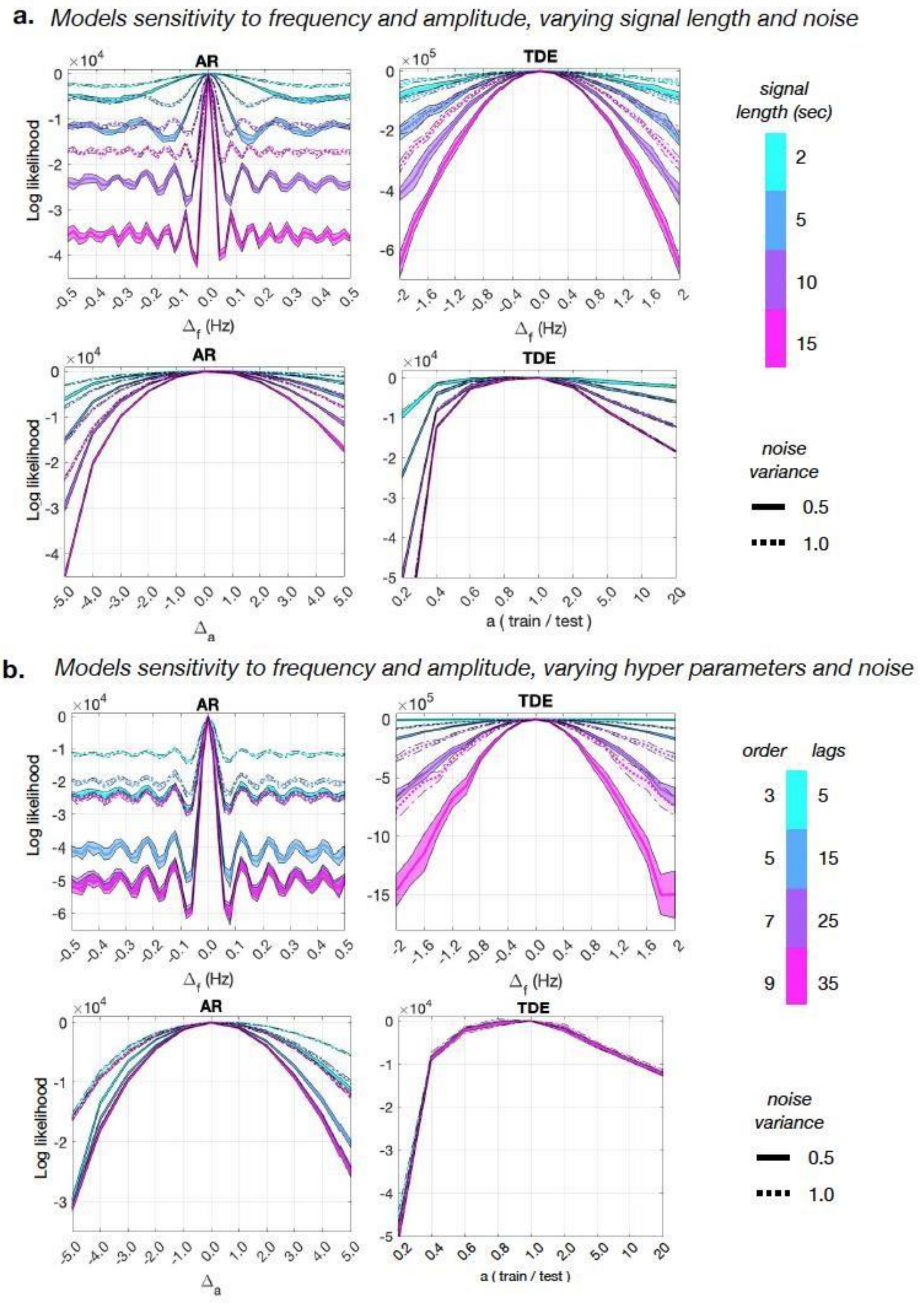


**Supplementary Figure 1:** Sensitivity of the observation models (AR and TDE) to frequency and amplitude as a function of signal length (amount of training data), models hyperparameters and noise, using *synthetic, stationary, one-channel data*. **a.** The plots show how the AR (left) and the TDE (right) models can tell apart two signals that differ only in frequency (top panels), or in amplitude (bottom panels), for different values of signal length and noise variance. Hyperparameters settings: $P$=3, $L$=21, $S$=1. **b.** AR (left) and the TDE (right) sensitivity to frequency (top panels), and amplitude (bottom panels) for different values of the models hyperparameters (order $P$ for the AR and lags $L$ for the TDE) and of the signals' noise variance. The TDE lags used here are all spaced in steps of $S$=1; signal length set to $T$=10 seconds.

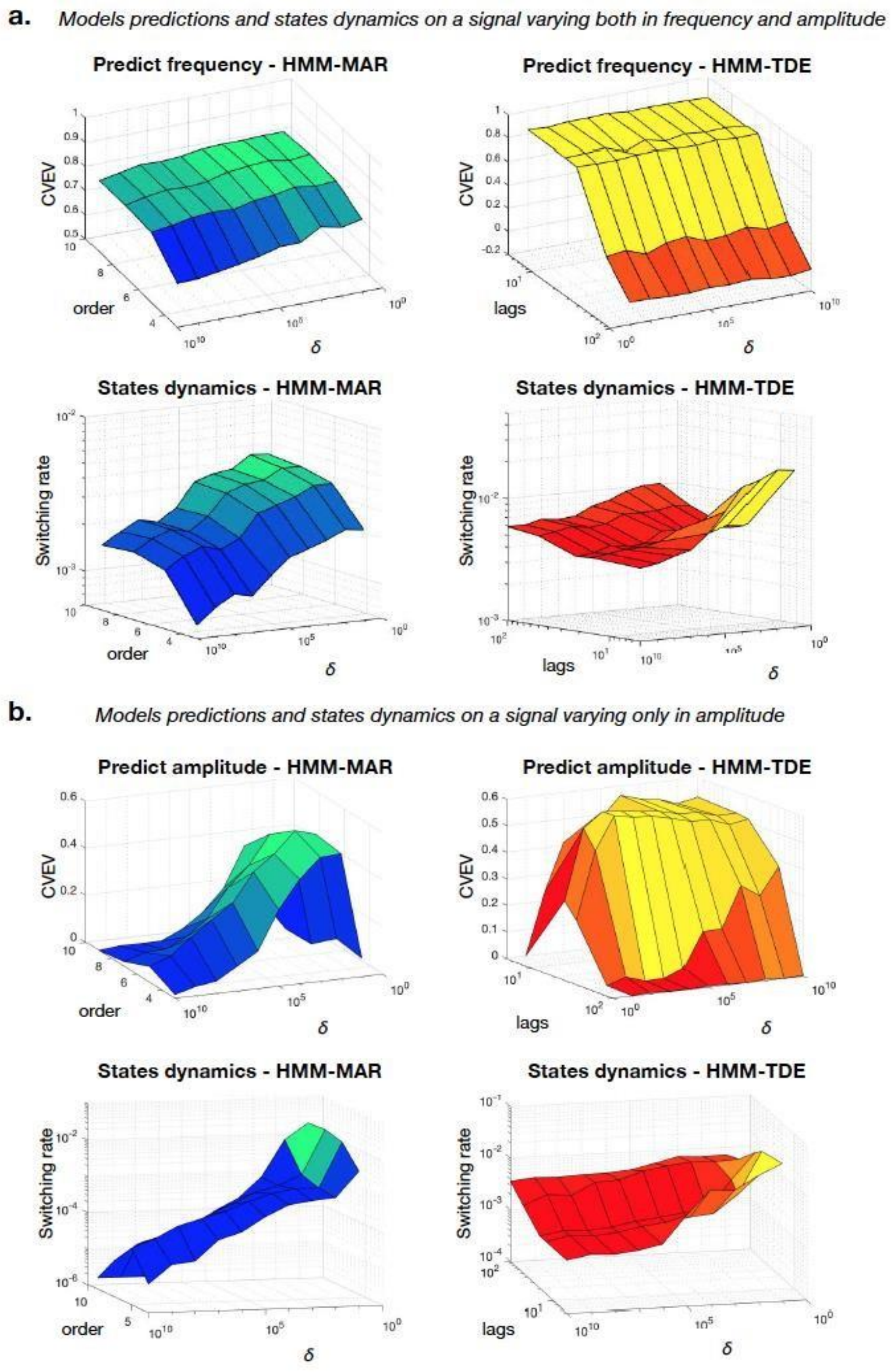


**Supplementary Figure 2:** Models prediction accuracy of frequency and amplitude (of *non-stationary, one channel signals*) as function of hyperparameters and *δ*, and the influence of *δ* on the states switching rate. **a.** **Top panels**: Accuracy of HMM-MAR (left) and HMM-TDE (right) states predicting the frequency of a signal varying both in frequency and amplitude, as a function of *δ* and the model specific hyperparameters; **bottom panels:** the states switching rate as a function of *δ* and the model specific hyperparameters for this analysis. Here, the HMM-TDE lags have values *L*=5,15,21,50,100, respectively in steps of *S*=1,3,4,5,10. Both the CVEV and the states switching rate are averaged over 20 repetitions of the experiment. Extension of **Figure 3c. b.** Same analysis and same settings described in **a**, applied to a signal varying only in amplitude. Extension of **Figure 3f**.

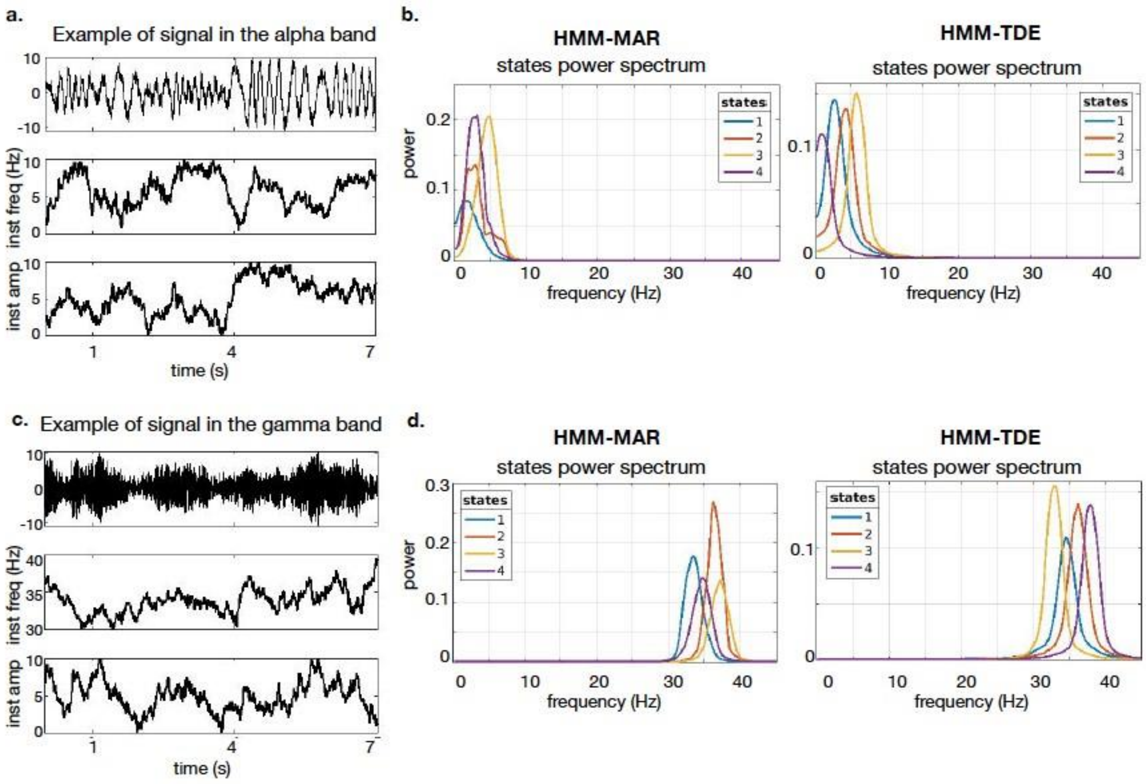


**Supplementary Figure 3**: Exploring HMM-MAR and HMM-TDE behavior on selected frequency bands. **a.** Example of signal in the alpha band. **b.** HMM-MAR (left panel) and HMM-TDE (right panel) states power spectra. **c.** Example of signal in the gamma band. **d.** HMM-MAR (left panel) and HMM-TDE (right panel) state power spectra. For these analyses, number of states $K$=4, $\delta$=10k, HMM-MAR order $P$=3, HMM-TDE lags $L$=15, $S$=3.

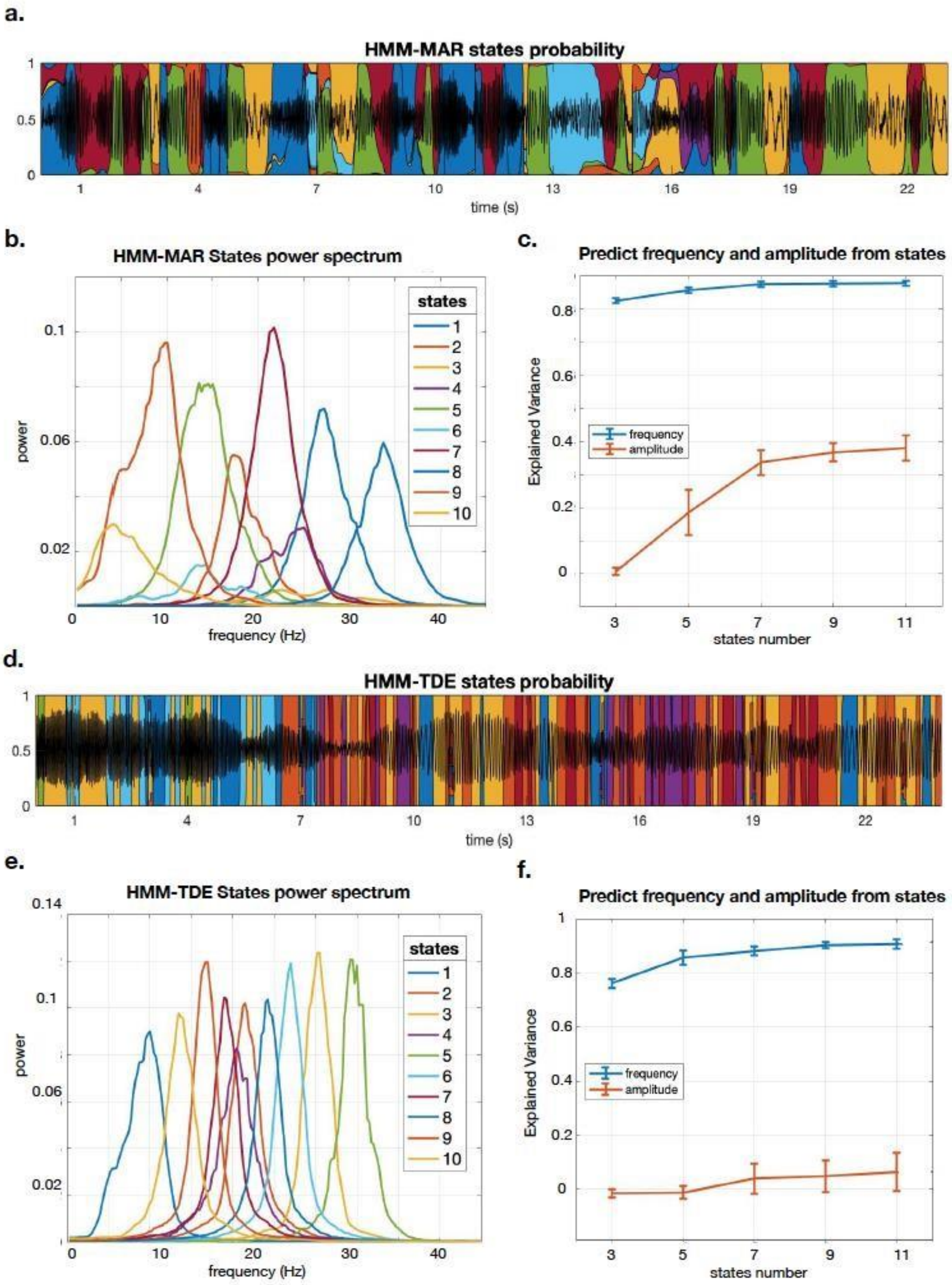


**Supplementary Figure 4**: Exploring the behavior of HMM-MAR and HMM-TDE with increasing number of states. **a.** Example of HMM-MAR state time courses. For this analysis, $P$=3, number of states $K$=10, $\delta$=1k. **b.** HMM-MAR state power spectra. **c.** Explained Variance of the regression analysis: HMM-MAR state time courses predicting instantaneous frequency and amplitude of the data, as a function of the number of states. **d.** Example of HMM-TDE state time courses; here, $L$=15, $S$=1, $K$=10, $\delta$=1k. **e.** HMM-TDE state power spectra. **f.** Same as in **c**, for HMM-TDE.

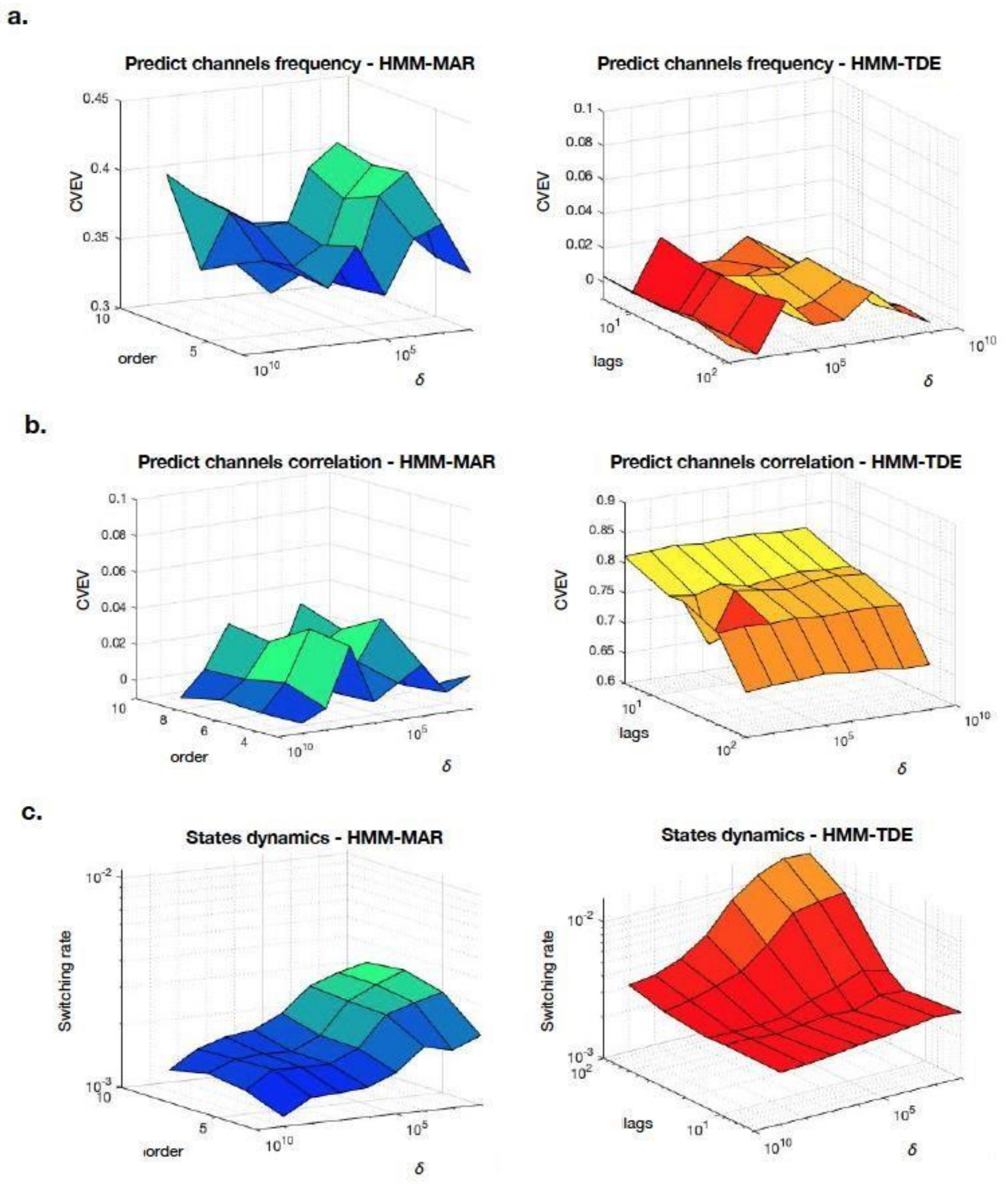


**Supplementary Figure 5:** Models prediction accuracy of channels frequency and between-channels correlation as function of hyperparameters and *δ*, and the influence of the *δ* on the states switching rate. **a.** Accuracy of HMM-MAR (left) and HMM-TDE (right) states predicting the channels frequency (average on the 2 channels), measured as Cross Validated Explained Variance (CVEV), as a function of *δ* and the model specific hyperparameters (order and lags, respectively). Extension of **Figure 4c** (left). **b.** Similar to **a**, prediction accuracy of between-channels correlation, extension of **Figure 4c** (right). **c.** The states switching rate for this analysis, for HMM-MAR (left) and HMM-TDE (right), as a function of δ and respective hyperparameters. Here, the HMM-TDE lags have values *L*=5,15,21,50,100, respectively in steps of *S*=1,3,4,5,10. Both the CVEV and the states switching rate are averaged over 10 repetitions of the experiment.

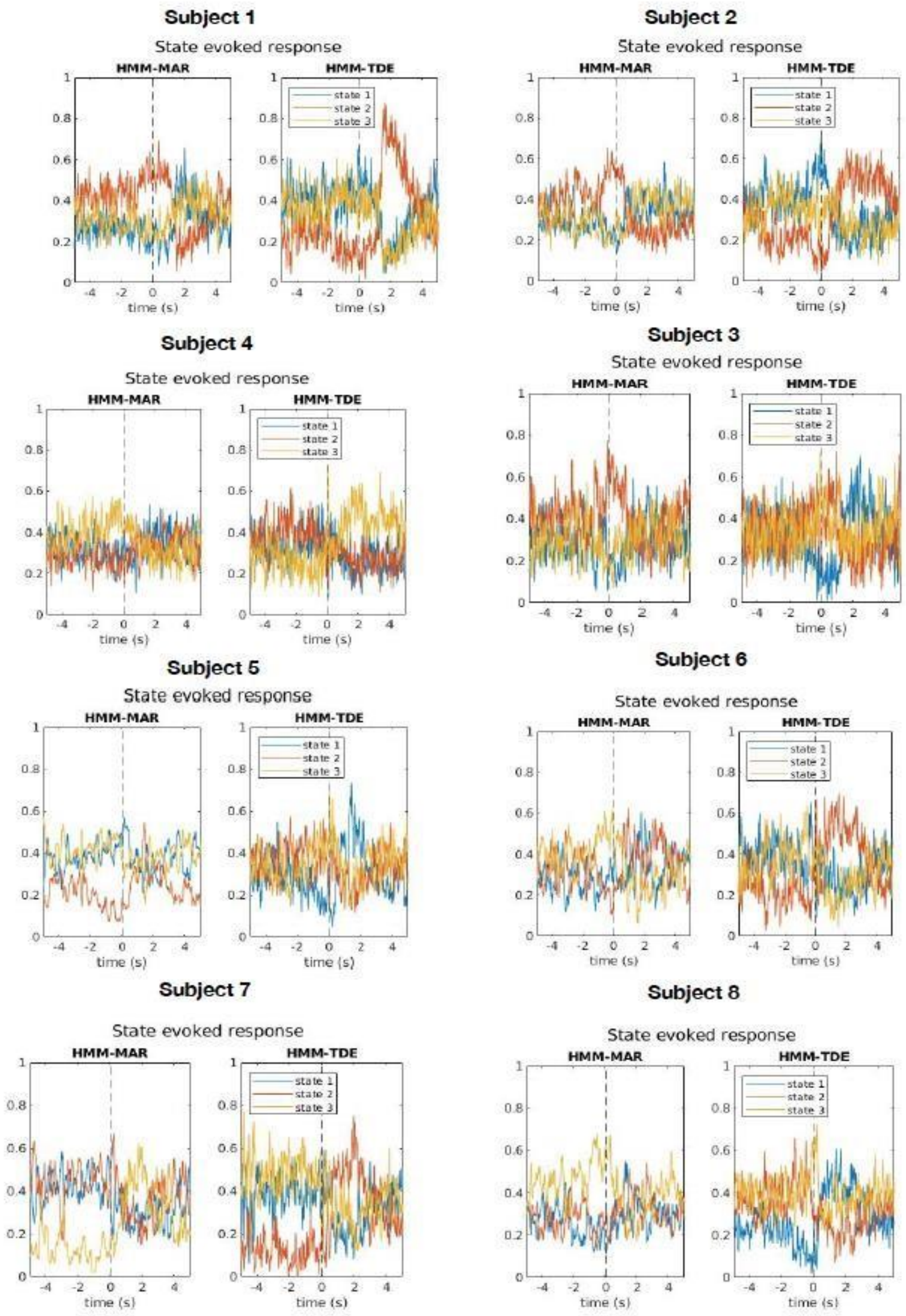


**Supplementary Figure 6**: The HMM state evoked responses of the complete set of subjects (8) for the MEG dataset analyzed. Completion of **Figure 7**, consistent with the results in Vidaurre *et al.* (2016).

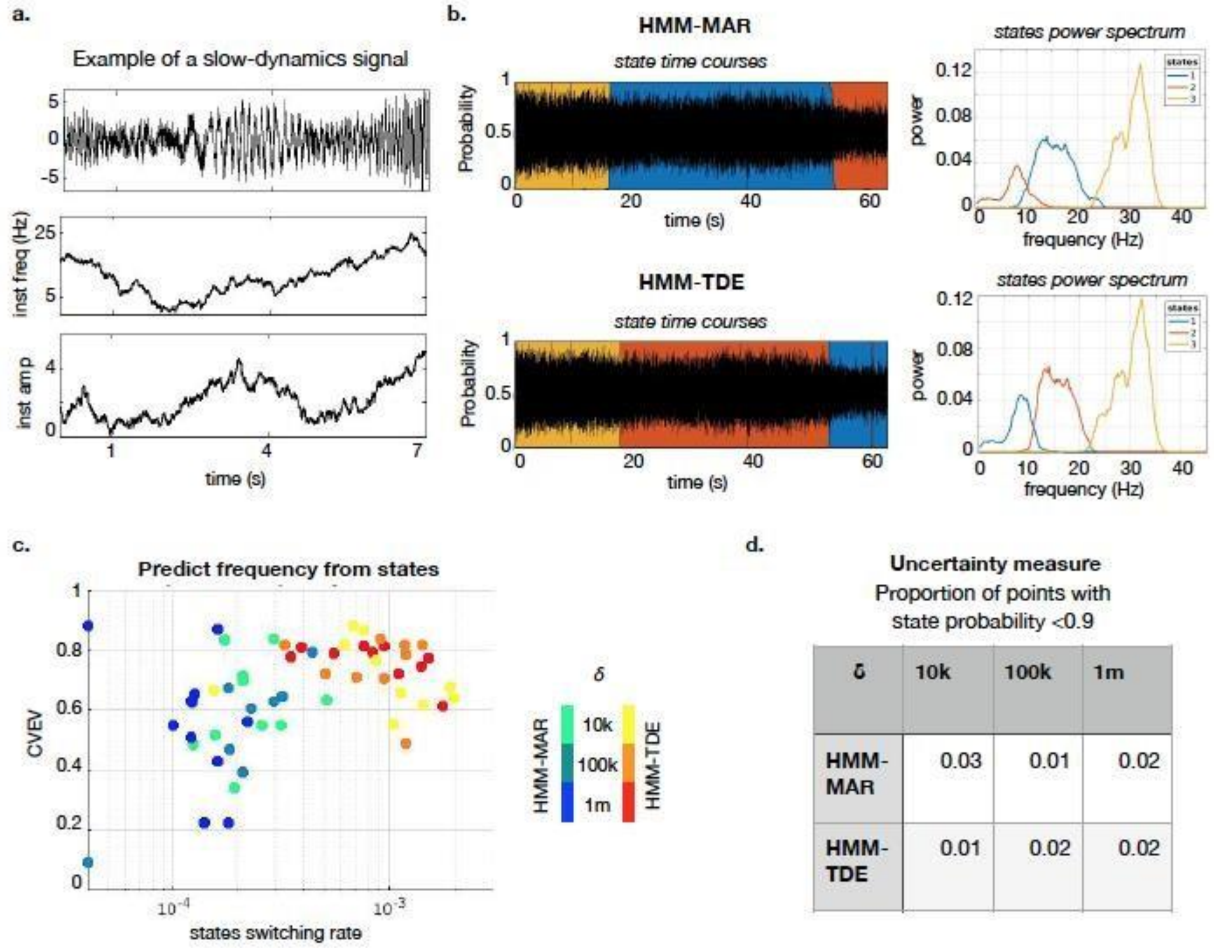


| δ | 10k | 100k | 1m |
|---|---|---|---|
| HMM-MAR | 0.03 | 0.01 | 0.02 |
| HMM-TDE | 0.01 | 0.02 | 0.02 |

**Supplementary Figure 7**: Exploring uncertainty in the HMM inference. **a.** Example of slowly varying signal. **b.** HMM-MAR (upper row) and HMM-TDE (lower row) states time courses and power spectra when applied to signal in **a.** **c.** Predicted frequency from the state time courses for different δ values. **d.** Uncertainty measure for slowly varying signal, average over 10 repetitions of each analysis scenario. For all the analyses, number of states $K$=3, HMM-MAR order $P$=3, HMM-TDE lags $L$=15, $S$=3.
